# Systematic Analysis of Mobile Genetic Elements Mediating β-lactamase Gene Amplification in Non-Carbapenemase-Producing Carbapenem Resistant *Enterobacterales* Bloodstream Infections

**DOI:** 10.1101/2022.05.20.492874

**Authors:** WC Shropshire, A Konovalova, P McDaneld, M Gohel, B Strope, P Sahasrabhojane, CN Tran, D Greenberg, J Kim, X Zhan, S Aitken, M Bhatti, TC Savidge, TJ Treangen, BM Hanson, CA Arias, SA Shelburne

## Abstract

Non-carbapenemase-producing carbapenem resistant *Enterobacterales* (non-CP-CRE) are increasingly recognized as important contributors to prevalent carbapenem resistant *Enterobacterales* (CRE) infections. However, there is limited understanding of mechanisms underlying non-CP-CRE causing invasive disease. Long- and short-read whole genome sequencing (WGS) was used to elucidate carbapenem non-susceptibility determinants in *Enterobacterales* bloodstream isolates at MD Anderson Cancer Center in Houston, Texas. We investigated carbapenem non-susceptible *Enterobacterales* (CNSE) mechanisms through a combination of phylogenetic analysis, antimicrobial resistant (AMR) gene detection/copy number quantification, porin assessment, and mobile genetic element (MGE) characterization. Most CNSE isolates sequenced were non-CP-CRE (41/79; 51.9%) whereas 25.3% (20/79) were carbapenem intermediate *Enterobacterales* (CIE) and 22.8% (18/79) were carbapenemase producing *Enterobacterales* (CPE). Statistically significant copy number variants (CNVs) of extended-spectrum β-lactamase (ESBL) genes (Wilcoxon Test; p-value < 0.001) were present in both non-CP-CR *E. coli* (median CNV = 2.6X; n= 17) and *K. pneumoniae* (median CNV = 3.2X, n = 17). All non-CP-CR *E. coli* and *K. pneumoniae* had predicted reduced expression of at least one outer membrane porin gene (*i.e., ompC/ompF* or *ompK36/ompK35*). Completely resolved CNSE genomes revealed that IS*26* and IS*Ecp1* structures harboring *bla*_CTX-M_ variants along with other AMR elements were the primary drivers of gene amplification, occurring in mostly IncFIB/IncFII plasmid contexts. MGE mediated β-lactamase gene amplifications resulted in either tandem arrays, primarily mediated by IS*26* ‘translocatable units’, or segmental duplication, typically due to IS*Ecp1* ‘transposition units’. Non-CP-CRE strains were the most prevalent cause of CRE bacteremia with carbapenem non-susceptibility driven by concurrent porin loss and MGE-mediated amplification of *bla*_CTX-M_ genes.

**IMPORTANCE:** Carbapenem resistant *Enterobacterales* (CRE) are considered urgent antimicrobial resistance (AMR) threats. The vast majority of CRE research has focused on carbapenemase producing *Enterobacterales* (CPE) even though non-carbapenemase-producing CRE (non-CP-CRE) comprise 50% or more of isolates in some surveillance studies. Thus, carbapenem resistance mechanisms in non-CP-CRE remain poorly characterized. To address this problem, we applied a combination of short- and long-read sequencing technologies to a cohort of CRE bacteremia isolates and used these data to unravel complex mobile genetic element structures mediating β- lactamase gene amplification. By generating complete genomes of 65 carbapenem non-susceptible *Enterobacterales* (CNSE) covering a genetically diverse array of isolates, our findings both generate novel insights into how non-CP-CRE overcome carbapenem treatments and provide researchers scaffolds for characterization of their own non-CP-CRE isolates. Improved recognition of mechanisms driving development of non-CP-CRE could assist with design and implementation of future strategies to mitigate the impact of these increasingly recognized AMR pathogens.

## INTRODUCTION

Carbapenem resistant *Enterobacterales* (CRE) infections are major public health challenges, particularly within vulnerable patient populations (1–6). There is a strong correlation between carbapenem and multi-drug resistance (MDR), in part because carbapenem resistant infections commonly occur in patients who have previously received multiple courses of antimicrobials (7, 8). A primary factor responsible for the dissemination of MDR phenotypes are mobile genetic elements (MGEs). These complex genetic structures (*e.g.*, plasmids, transposons, integrons) can mobilize carbapenem resistance determinants in addition to other antimicrobial resistance (AMR) genes that confer resistance to other classes of antibiotics such as fluoroquinolones, aminoglycosides, and other novel β-lactam/β-lactamase inhibitor combinations (9–13). In recent years, the development of long-read sequencing technologies has improved our understanding of the complexity, diversity, and prevalence of these MGEs as key drivers of MDR infections (13–18).

There are two general mechanisms by which MGEs contribute to the development of carbapenem resistance in *Enterobacterales* (19). MGEs can disseminate and mobilize carbapenemases, enzymes that are able to hydrolyze the carbapenem β-lactam ring with sufficient efficiency to inactivate the drug, through horizontal gene transfer pathways (11, 20). For example, there are well documented associations of the *Klebsiella pneumoniae* carbapenemase (KPC) encoding gene being disseminated through isoforms of the Tn3-based Tn*4401* transposon (21). Interestingly, in recent years, surveillance studies have found that up to 50% of CRE detected lack a carbapenemase, *i.e.*, are non-carbapenemase producing CRE (non-CP-CRE) (1–3). Similar to MGEs key role in dissemination of carbapenemases, MGEs are also necessary for the dissemination of extended-spectrum β-lactamase (ESBL) and AmpC-like encoding enzymes that are both critical for the development of the non-CP-CRE phenotype (11, 12, 22–26).

Much of the existing knowledge regarding non-CP-CRE mechanisms is derived from laboratory passaging or serial, single isolate studies (22–26). These studies have shown that non-CP-CRE development typically involves increased expression or gene copy number of ESBL or AmpC-like enzymes in conjunction with outer membrane porin (*omp*) gene inactivation, which results in a reduced carbapenem concentration in the periplasmic space (22–26). Given that both ESBL and AmpC-like encoding genes are typically located in MGEs (11, 13, 27), an increase in β-lactamase gene copy number would seem to be feasible for a broad array of ESBL and AmpC-like positive *Enterobacterales*. To our knowledge, a systematic analysis of MGE mediated β-lactamase encoding gene amplifications in a large cohort of CRE isolates using completed genome assemblies has not been performed. Given the repetitive, complex nature of MGEs that harbor these β-lactamase encoding genes, PCR detection or short-read sequencing approaches have had limited capacity to reveal the breadth of MGEs contributing to these varied CRE phenotypes.

Herein, we sought to systematically determine carbapenem resistance mechanisms by applying a combination of short- and long-read sequencing to a well-defined cohort of carbapenem non-susceptible *Enterobacterales* (CNSE) isolates. We found that non-CP-CRE isolates caused the vast majority of our CRE bacteremia cases and harbored MGEs with complex arrangements primarily of ESBLs, such as *bla*_CTX-M_ variants, mediated by either IS*26* or IS*Ecp1* elements.

There was a statistically significant association of ESBL amplification in conjunction with *omp* gene disruption in non-CP-CR *Escherichia coli* and *Klebsiella pneumoniae.* Using Oxford Nanopore Technologies (ONT) long-read sequencing, we clarified that ESBL amplification was caused by IS*26*-mediated ‘translocatable units’ (TUs) and IS*Ecp1* ‘transposition units’ (TPUs) in both non-CP-CR *Escherichia coli* and *Klebsiella pneumoniae* thereby improving the understanding of mechanisms underlying the non-CP-CRE phenotype.

## RESULTS

### Molecular Epidemiology of Carbapenem Non-Susceptible *Enterobacterales* (CNSE) Causing Bacteremia at MD Anderson Cancer Center (MDACC)

There were 1632 unique *Enterobacterales* BSIs at our institution from July 2016 to June 2020. The leading causes were *Escherichia coli* (939/1632; 57.5%) followed by *Klebsiella pneumoniae* (338/1632; 20.7%) and *Enterobacter spp.* (159/1632; 9.7%). A total of 5.2% (85/1632) were CDC-defined carbapenem resistant with an additional 1.8% (29/1632) having intermediate carbapenem resistance based on CLSI guidelines (*i.e.*, carbapenem intermediate *Enterobacterales* [CIE]), resulting in a total 7.0% (114/1632) that were carbapenem non-susceptible *Enterobacterales* (CNSE) as initially determined by the MDACC clinical microbiology laboratory. When stratifying the causal species of BSI by carbapenem non-susceptibility, 39.5% (45/114) of CNSE were *Escherichia coli* followed by *Klebsiella pneumoniae sensu stricto* (30.7%; 35/114) and *Enterobacter spp.* (16.7%; 19/114). We found a statistically significant difference in carbapenem non-susceptibility by species (Fisher’s exact test p-value < 0.001) with a higher prevalence of *K. pneumoniae* BSIs (10.4%; 35/338) that were carbapenem non-susceptible as compared to *E. coli* (4.8%; 45/939), consistent with other CRE surveillance studies in the United States (1, 2, 28).

A total of 91% (104/114) CNSE BSI isolates were present in our sample collection **(Figure 1)**. Of these 104 CNSE BSI isolates, we confirmed *at least* ertapenem MIC intermediate interpretations for 37/42 *E. coli* (88%), 28/32 *K. pneumoniae* (88%), 8/15 *Enterobacter spp.* (53%), and 6/15 other *Enterobacterales* (40%) with the remaining isolates being considered unconfirmed-CNSE **(Figure 1)**. Thus, we had 79 CNSE-confirmed BSI isolates that we could perform whole genome sequencing (WGS) on to determine respective carbapenem non-susceptibility genotypes. Only 23% of BSI isolates (18/79) had a confirmed carbapenemase whereas the majority were non-CP-CRE (41/79; 52%) or CIE (20/79; 25%) based on WGS analysis and carbapenem MIC determination **(Figure 1)**. We identified 17 CNS*Ec* bacteremia cases that had a prior initial carbapenem susceptible *E. coli* bacteremia infection which had tested positive for ESBL production in 16/17 cases. Interestingly, all 17 of these CNS*Ec* isolates were carbapenemase negative. Similarly, 5/6 CNS*Kp* that were preceded by an initial carbapenem susceptible *K. pneumoniae* bacteremia were carbapenemase negative as well. When focusing on clinical features, there were no statistically significant differences in age, gender, country of origin, recent travel history, or predicted source of BSI across each of the CNSE categories, albeit there were a small number of observations per category (Table S1).

**Figure 1:**
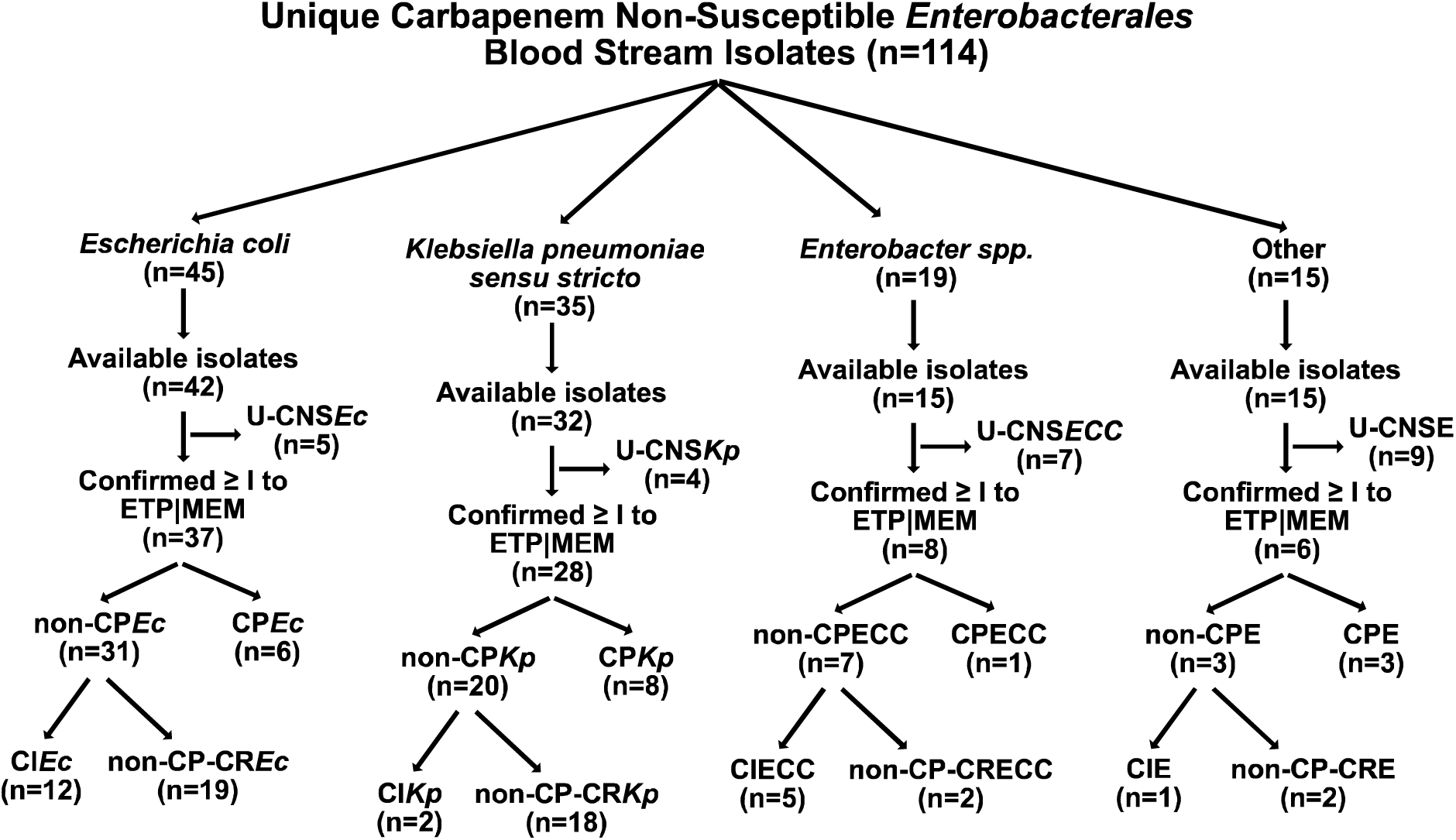
Selection and Delineation of Carbapenem Non-Susceptible *Enterobacterales* Blood Stream Infection Isolates. Total isolates per group included in parenthesis. U-CNS = unconfirmed carbapenem non-susceptible; non-CP = non-carbapenemase producing; non-CP-CR = non-carbapenemase-producing carbapenem resistant; CP = carbapenemase producing; *Ec* = *Escherichia coli*; *Kp* = *Klebsiella pneumoniae*; ECC = *Enterobacter cloacae* complex; E = *Enterobacterales*.

*Enterobacter spp.* were the third most prevalent group of CNSE BSI isolates with all isolates belonging to the *Enterobacter cloacae* complex (ECC) (Table S2). The majority of CNSE-confirmed ECC had CIE phenotypes (5/8; 63%), with only one carbapenemase-producing ECC (CPECC) isolate harboring *bla*_KPC-2_ (MB8139), and two non-carbapenemase-producing carbapenem resistant ECC (non-CP-CRECC). With regards to the non-CP-CRECC isolates, both had outer membrane porin (*omp*) gene disruptions with one non-CP-CRECC (MB5921) containing an ESBL gene (*bla*_SHV-12_). The other non-CP-CRECC isolate (MB6956) had a carbapenem resistant mechanism that likely involved an overexpressed chromosomal *ampC* gene (*bla*_CMH_) due to an *ampD*/*ampE* fusion mutation, with the inactivation of the AmpD gene predicted to result in AmpC derepression (29) (Table S2). The six other *Enterobacterales* spp. detected in our cohort included 3 CPE (*Klebsiella spp.* not including *K. pneumoniae sensu stricto*), 2 non-CP-CRE (1 *K. aerogenes,* 1 *Citrobacter freundii*), and 1 CIE (*Serratia marcescens*) (Table S2). We focused the remainder of this study on the two most common, clinically relevant species in our cohort, *E. coli* and *K. pneumoniae*, and the putative mechanisms responsible for their carbapenem non-susceptible phenotypes.

### Characterization of carbapenem resistance mechanisms among CNS *E. coli* and *K. pneumoniae* isolates

There were 37 unique carbapenem non-susceptible *E. coli* (CNS*Ec*) bacteremia isolates with 6 CP*Ec* (16%), 19 non-CP-CR*Ec* (51%), and 12 CI*Ec* (32%) (Table S2). Core gene alignment inferred, maximum-likelihood phylogenetic trees for CNS*Ec* isolates with carbapenem susceptibility profile, outer membrane porin gene (*omp*) mutation status, and β-lactamase gene presence/absence with copy number estimates are shown in **Figure 2A**. Hierarchical clustering of core gene SNPs resulted in five clusters, indicated by tip label color **(Figure 2A)**, that segregate isolates based on phylogroups A (n=12), B2 (n=11), D (n=7), B1/C (n=8), and F (n=2) (30). The most identified sequence type (ST) among CNS*Ec* was the pandemic, uropathogenic strain ST131 (10/37; 27%). The mean pairwise core gene SNP difference was 57355 SNPs (S.D. = 25621 SNPs). There were only two clinical isolates, MB9272 and MB9880, that had less than 50 core gene SNP differences (18 SNPs), further indicating minimal clonal infections amongst the *E. coli* strains in our cohort.

**Figure 2:**
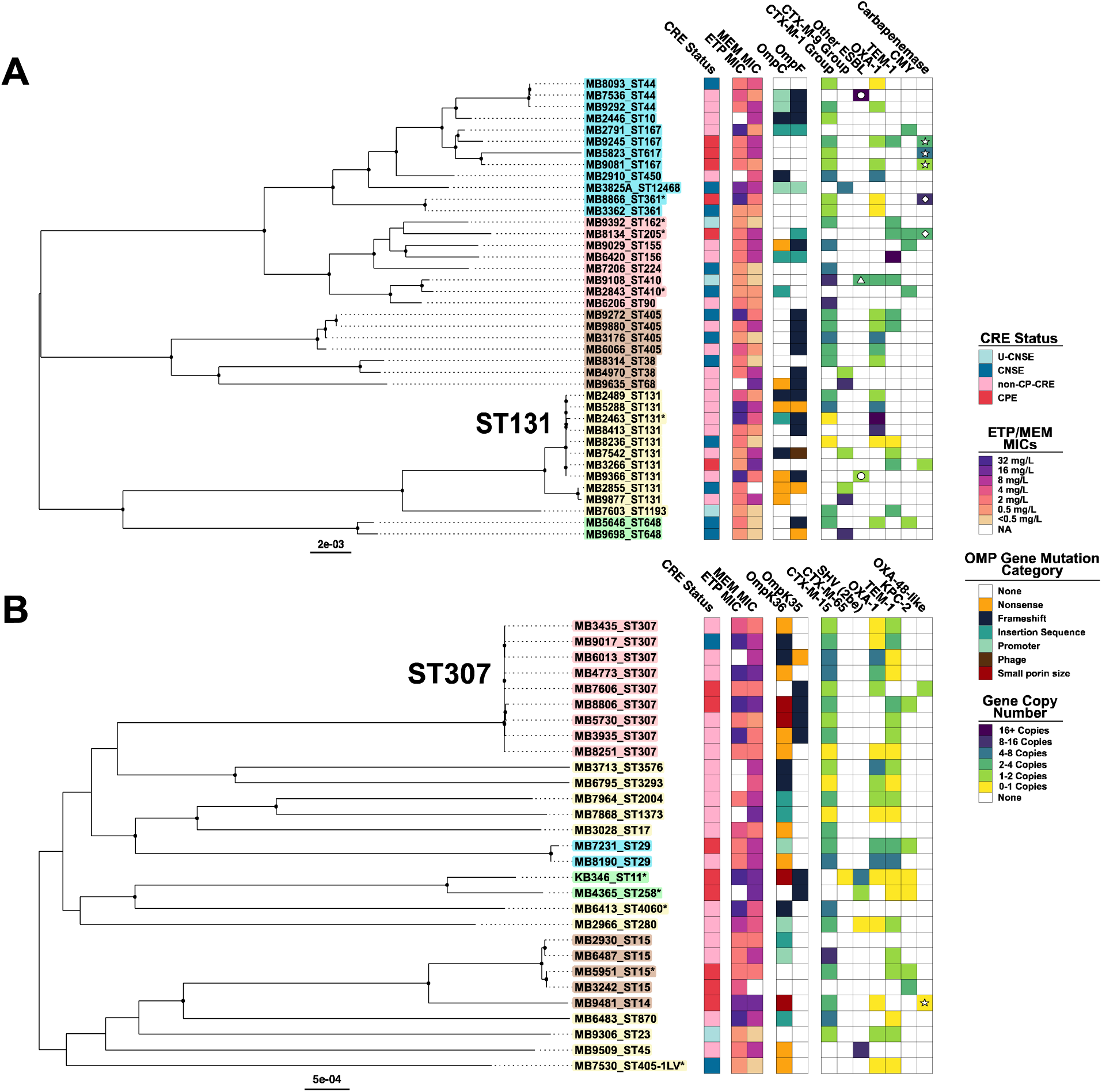
Population structure of *E. coli* and *K. pneumoniae* bacteremia isolates with phenotype/genotype data. Core gene alignment inferred, mid-point rooted maximum likelihood phylogenies. Circles at internal nodes indicate UFBoot values with ≥95% support. Tip label background color corresponds to nested population structure identified using hierarchical clustering of sequence data with rhierbaps. Carbapenem resistance status, ertapenem (ETP) and meropenem (MEM) MICs (μg/mL), outer membrane porin gene mutation status, and gene copy number estimate are presented in columnar data from left to right and labelled in the legend respectively. ‘*’ adjacent to tip label indicates isolates with only draft assembly. **(A)** *E. coli* population structure. Circles in ‘Other ESBL’ column indicate *bla*_TEM_ variants whereas triangle indicates *bla*_SHV-12_. Stars in ‘carbapenemase’ indicate *bla*_NDM-5_, diamond indicates *bla*_OXA-48-like_, and absence of shape indicates *bla*_KPC-2_. **(B)** *K. pneumoniae* population structure. Isolate with star in ‘*bla*_OXA-48-like_’ column indicates co-carriage of *bla*_NDM-1_ with 1-2 copies.

Among the six CP*Ec* isolates, three isolates from phylogroup A harbored *bla*_NDM-5_, two unique ST isolates harbored plasmid borne *bla*_OXA-48-like_ genes (MB8866 = *bla*_OXA-232_; MB8134 = *bla*_OXA-181_), and one isolate (MB3266) carried a plasmid-borne Tn*4401a* transposon harboring *bla*_KPC-2_.

Only one CP*Ec* (MB8134) had an *omp* mutation (IS*2* insertion within *ompF*) **(Figure 2A)**. With regards to non-CP-CR*Ec*, 79% (15/19) of isolates were ESBL positive. The most common β-lactamases detected in non-CP-CR*Ec* were CTX-M-1 group variants (7 *bla*_CTX-M-15_; 3 *bla*_CTX-M-55_), CTX-M-9 group variants (3 *bla*_CTX-M-27_; 1 *bla*_CTX-M-14_; 1 *bla*_CTX-M-195_), *bla*_OXA-1_ (n=8), *bla*_TEM-1_ (n=4), and *bla*_CMY_ variants (n=2). One ST131 non-CP-CR*Ec* isolate (MB9366) carried a novel *bla*_TEM_ variant (p.M182T, p.G238S, p.E240K, p.S243A, p.S270G), which was identified as an ESBL-E by the MDACC clinical microbiology lab and has an antibiogram that resembles an ESBL-E (Table S3). In contrast to the low prevalence of *ompC* and *ompF* mutations detected in CP*Ec*, all 19 non-CP-CR*Ec* isolates had at least one *ompC* or *ompF* mutation except for MB6206 **(Figure 2A)**, which had an IS*Ecp1*-*bla*_CTX-M-55_ insertion into the histidine kinase gene *envZ*, a known regulator of *ompC* and *ompF* expression (31). Consistent with EnvZ inactivation, immunoblot analysis confirmed a significant reduction of OmpC/OmpF in MB6206 (Supplemental Figure 1). Furthermore, 63% (12/19) of non-CP-CR*Ec* isolates were double mutant *ompC/ompF* isolates **(Figure 2A)**. Similar to non-CP-CR*Ec*, 11/12 (91.7%) of CI*Ec* were ESBL carriers with eight CTX-M-1 group variants (7 *bla*_CTX-M-15_;1 *bla*_CTX-M-1_) and three CTX-M-9 group variants (2 *bla*_CTX-M-14_;1 *bla*_CTX-M-27_). Other common β-lactamases detected in CI*Ec* were *bla*_OXA-1_ (n=7) and *bla*_CMY_ (n=2) variants. Relative to non-CP*Ec* (18/19), CI*Ec ompC* and *ompF* mutations were less prevalent (7/12; 58%; Fisher’s Exact Test p-value = 0.02) with only two strains (16%) having mutations in both genes.

There were 28 unique carbapenem non-susceptible *K. pneumoniae* (CNS*Kp)* bacteremia isolates with eight CP*Kp* (29%), 18 non-CP-CR*Kp* (64%), and two CI*Kp* (7%) (Table S2). The core population structure of CNS*Kp* BSI isolates is presented in **Figure 2B**. The finding that 64% CR*Kp* were non-carbapenemase producers was noteworthy given that in most US based CRE surveillance studies, the majority of CR*Kp* are carbapenemase positive (1, 28). Indeed, for our cohort the proportion of non-CP-CR*Kp* (18/28) was comparable to non-CP-CR*Ec* isolates (19/37; χ-squared test statistic = 0.62; p-value = 0.4). The most common sequence type identified was the ST307 lineage (9/28; 32%) followed by 18% (5/28) belonging to clonal group 15 (CG15). Hierarchical clustering demonstrated that, apart from ST307 and CG15 isolates, most CNS*Kp* belonged to single, long-branching isolates (**Figure 2B**), indicating limited genetic relatedness. In support of this observation was a mean pairwise core gene SNP difference of 22141 SNPs (S.D. 7864) with the minimum pairwise core gene SNP distance between our CNS*Kp* isolates being 38 SNPs between two ST15 isolates (MB5951 and MB3242). Amongst CP*Kp*, six isolates encoded *bla*_KPC-2_, one isolate (MB7606) encoded *bla*_OXA-181_, and one isolate (MB9481) encoded two carbapenemases, *bla*_NDM-1_ and *bla*_OXA-48_. *OmpK36* or *ompK35* mutations (*i.e., ompC* and *ompF K. pneumoniae* homologs respectively) that would be predicted to affect outer membrane porin function were present in 5/8 (62.5%) CP*Kp*. Almost all non-CP-CR*Kp* carried *bla*_CTX-M-15_ (16/18; 84%) with one such isolate having a novel, single amino acid *bla*_CTX-M-15_ variant (MB6013; p.P269S). The β-lactamase encoding genes *bla*_OXA-1_ (n = 14) and *bla*_TEM-1_ (n=10) were also commonly detected in non-CP-CR*Kp.* All non-CP-CR*Kp* isolates had an *ompK36* mutation with 16.6% (3/18) also having an *ompK35* mutation **(Figure 2B)**. Only 2 CI*Kp* isolates were identified, both having *ompK36* disrupted ORFs with one isolate (MB9017) harboring *bla*_CTX-M-15_, *bla*_OXA-1_, and *bla*_TEM-1_ and the other isolate harboring only *bla*_OXA-1_ and *bla*_TEM-1._ Taken together, the core population structure indicates disparate CNS *E. coli* and *K. pneumoniae* sequence types with little evidence of clonal outbreaks in addition to a high prevalence of ESBL encoding genes with universal predicted *omp* gene disruption within non-CP-CRE isolates.

### Copy number variant profiling of **β**-lactamases encoding genes in CNSE

An increase in copy number of ESBL, AmpC-like, and narrow-spectrum β-lactamase encoding genes has been previously documented as contributing to CNSE development (13, 23, 25, 26). Thus, we next sought to comprehensively assess the presence of β-lactamase gene amplifications and their associations with each carbapenem non-susceptibility profile (Table S4). To this end, we analyzed β-lactamase encoding gene copy number variants (CNVs) and determined which CNSE groups had median CNV estimates greater than baseline (*i.e.,* 1 copy) **(Figure 3)**. Non-CP-CR*Ec* contained statistically significant increases in gene copy numbers of the narrow spectrum β-lactamase encoding gene *bla*_OXA-1_ (median CNV = 3.4X; one-sample, one-sided, Wilcoxon Test p-value=0.004) **(Figure 3A)** that were not found in other CNS*Ec* categories nor in any of the CNS*Kp* groups **(Figure 3B)**. Both non-CP-CR*Ec* (median CNV = 2.6X; Wilcoxon Test p-value <0.0001) and non-CP-CR*Kp* (median CNV = 3.2X; Wilcoxon Test p-value <0.001) had statistically significant increases in ESBL gene copy numbers shown in **Figure 3C** and **Figure 3D** respectively. Notably 80% (12/15) and 64% (11/17) of ESBL positive, non-CP-CR*Ec* and non-CP-CR*Kp* respectively had an estimated ≥2 copies of ESBL encoding genes (Table S4). Similar to non-CP-CR*Ec*, CI*Ec* also had a statistically significant increase in ESBL gene copy number (median CNV = 2.6X; p-value <0.001). Amplification of carbapenemase encoding genes (median CNV = 2.4X; p-value = 0.02) was also detected in CP*Ec* **(Figure 3E)**, which was not evident in CP*Kp* (median CNV = 1.4X; p-value = 0.2) **(Figure 3F)**. While there was notably high *bla*_TEM-1b_ amplification in non-CP-CR*Ec* (median CNV = 11.5X), this did not reach statistical significance likely due to small number of observations (n=4) and high variance in CNV estimates (Supplemental Figure 2A); whereas non-CP-CR*Kp* did not have evidence of *bla*_TEM-1b_ amplification (Supplemental Figure 2B). Lastly, *bla*_CMY_ amplification was present in CNS*Ec* with all five *bla*_CMY_ positive isolates having estimated copy numbers greater than two (Table S4). Thus, a broad range of β-lactamases had evidence of gene copy number amplifications with statistically significant ESBL gene amplifications being detected in both non-CP-CR*Ec* and non-CP-CR*Kp* isolates.

**Figure 3.**
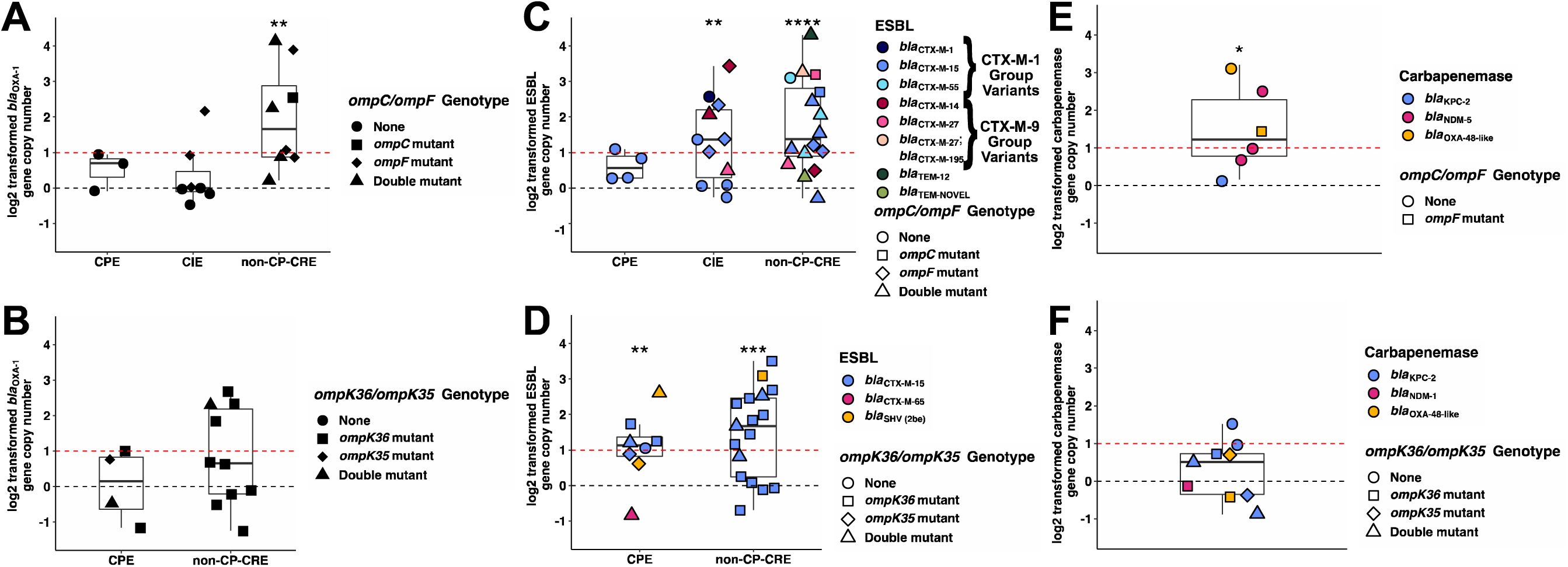
Log2 transformed β-lactamase gene copy numbers with outer membrane porin gene mutation profile stratified by carbapenem non-susceptible (CNS) definitions. (A,C,E) *Escherichia coli* and (B,D,F) *Klebsiella pneumoniae* CNS isolates. Black dotted horizontal line at y=0 is equivalent to 1X gene copy; Red dotted horizontal line at y=1 is equivalent to 2X gene copy. CPE = carbapenemase-producing *Enterobacterales*; CIE = carbapenem ‘intermediate’ *Enterobacterales*; non-CP-CRE = non-carbapenemase-producing carbapenem resistant *Enterobacterales.* One sample, one-sided, Wilcoxon Test on non-transformed copy number estimates to determine statistically significant gene copy number amplifications (*i.e.*, > 1 copy) with *P*-value: *<0.05; **<0.01; ***<0.001, ****<0.0001.

### Genomic structures contributing to carbapenem resistance development in CNSE cohort

Having quantified the extent of β-lactamase amplification across each of the CNSE groups, we used long-read, ONT sequencing to complete genomes of 65 CNSE isolates (Table S2) in order to resolve the putative MGEs responsible for mobilization and amplification of β-lactamase encoding genes. We initially characterized the MGEs in CNSE isolates harboring β-lactamase genes greater than or equal to 2X copies **(Figure 3)** with results shown for CNS*Ec* **(Table 1)** and CNS*Kp* **(Table 2)**. When we subset these isolates with complete genomes available, we found the majority of CNS*Ec* (21/27; 78%) and CNS*Kp* (12/15; 80%) had MGE *in situ* tandem or *ex situ* segmental duplication contributing to increased β-lactamase copy numbers. Furthermore, with rare exception, these β-lactamase amplifications were mediated by the widely observed insertion sequences IS*26* and/or IS*Ecp1* within CNSE genomes **(Table 1** and **Table 2)**. Stratifying by species and using nomenclature established for these aforementioned MGEs (27), for the 21 CNS*Ec* with MGE mediated β-lactamase gene amplification, 11 (52%) had IS*26* ‘translocatable units (TUs)’, eight (38%) had IS*Ecp1* ‘transposition units (TPUs)’, and one isolate had both mechanisms **(Table 1)**. Conversely, of the 12 CNS*Kp* with at least two copies of B-lactamase encoding genes driven by MGEs, eight (67%) had TPUs, three had (25%) TUs, and one isolate had both mechanisms **(Table 2)**. Thus, IS*26* mediated TU or IS*Ecp1* mediated TPU amplifications were the primary drivers of MGE inter- and intramolecular mobilization of β- lactamases that contributed to carbapenem non-susceptibility.

**Table 1:**
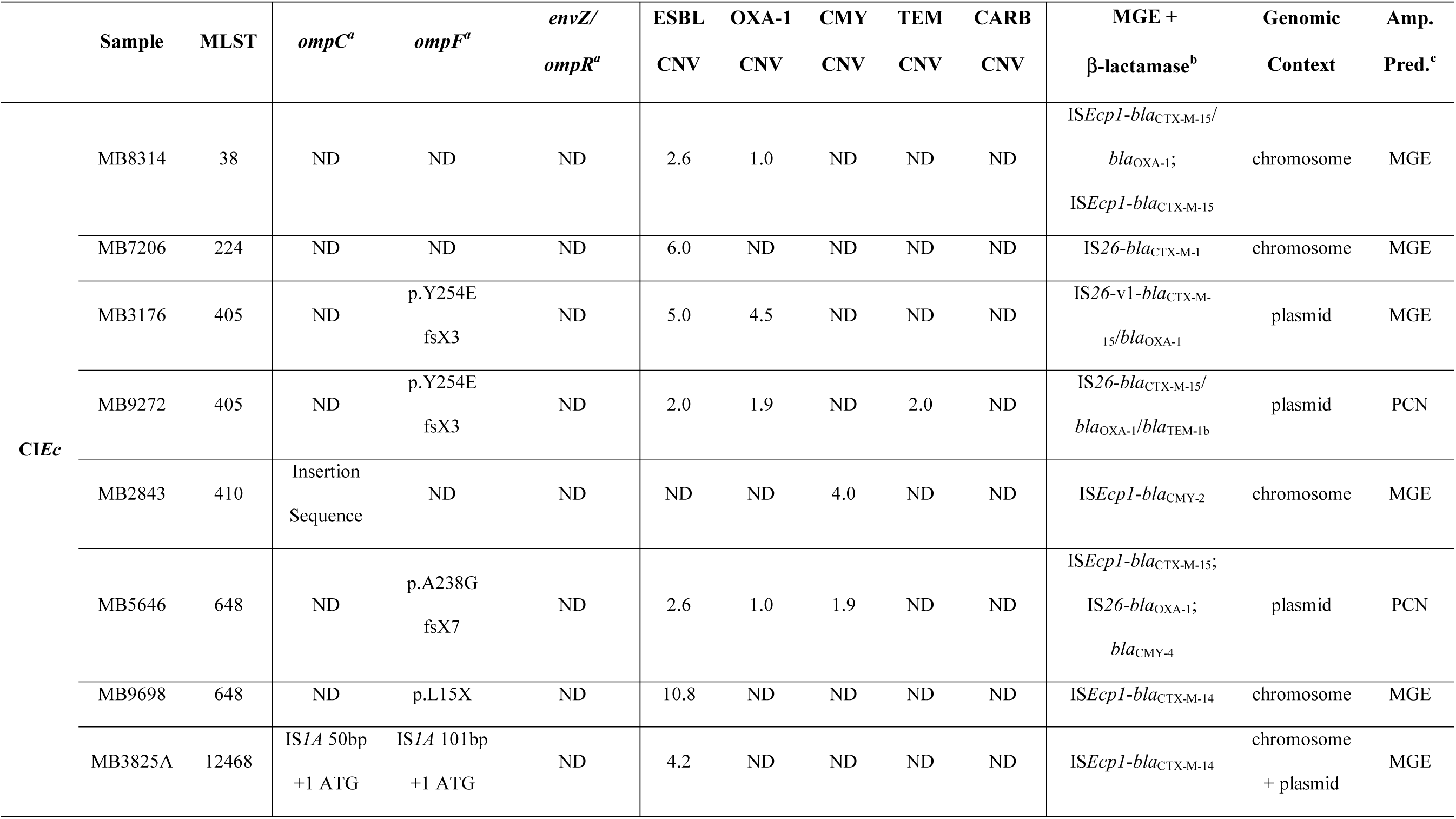
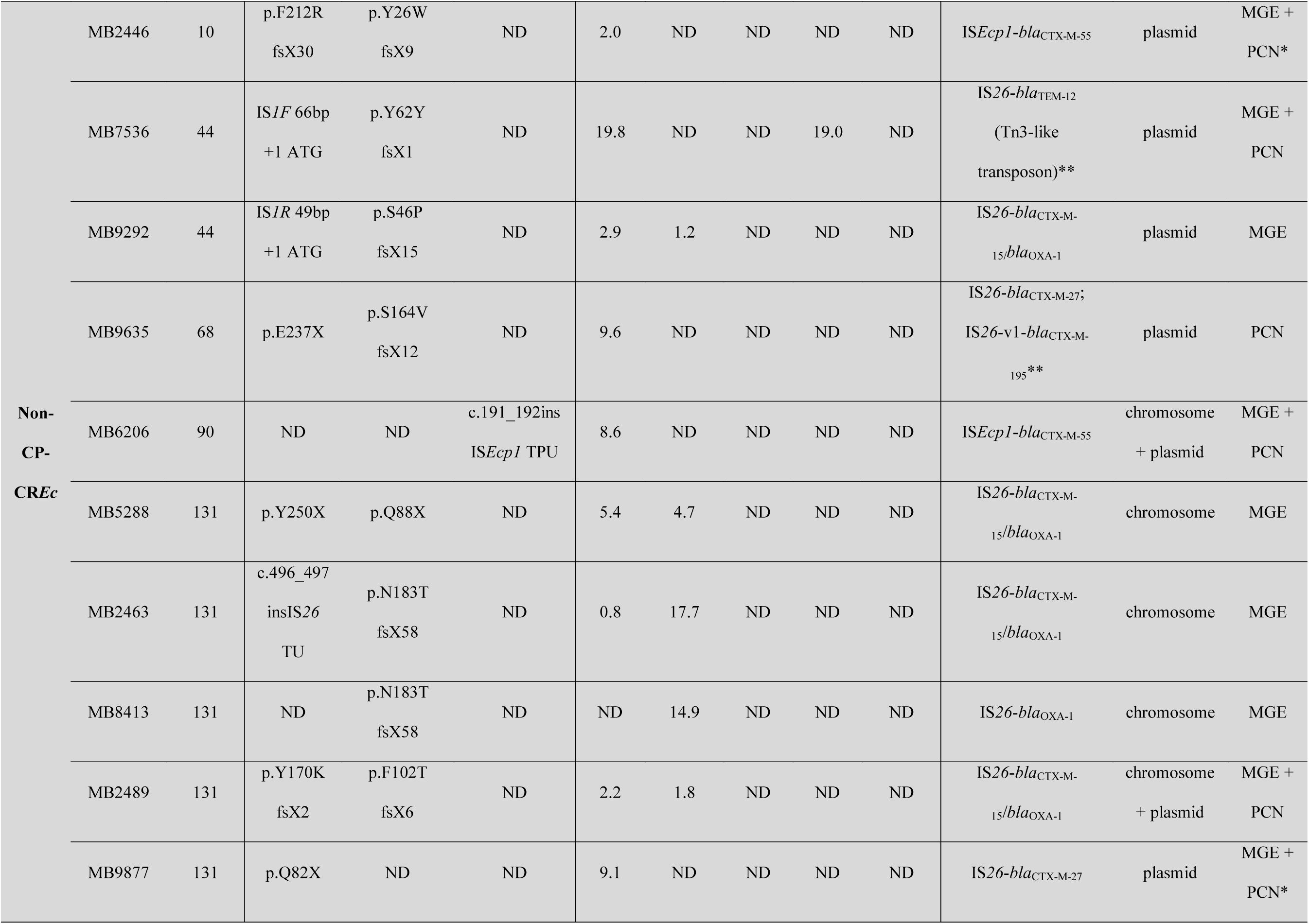
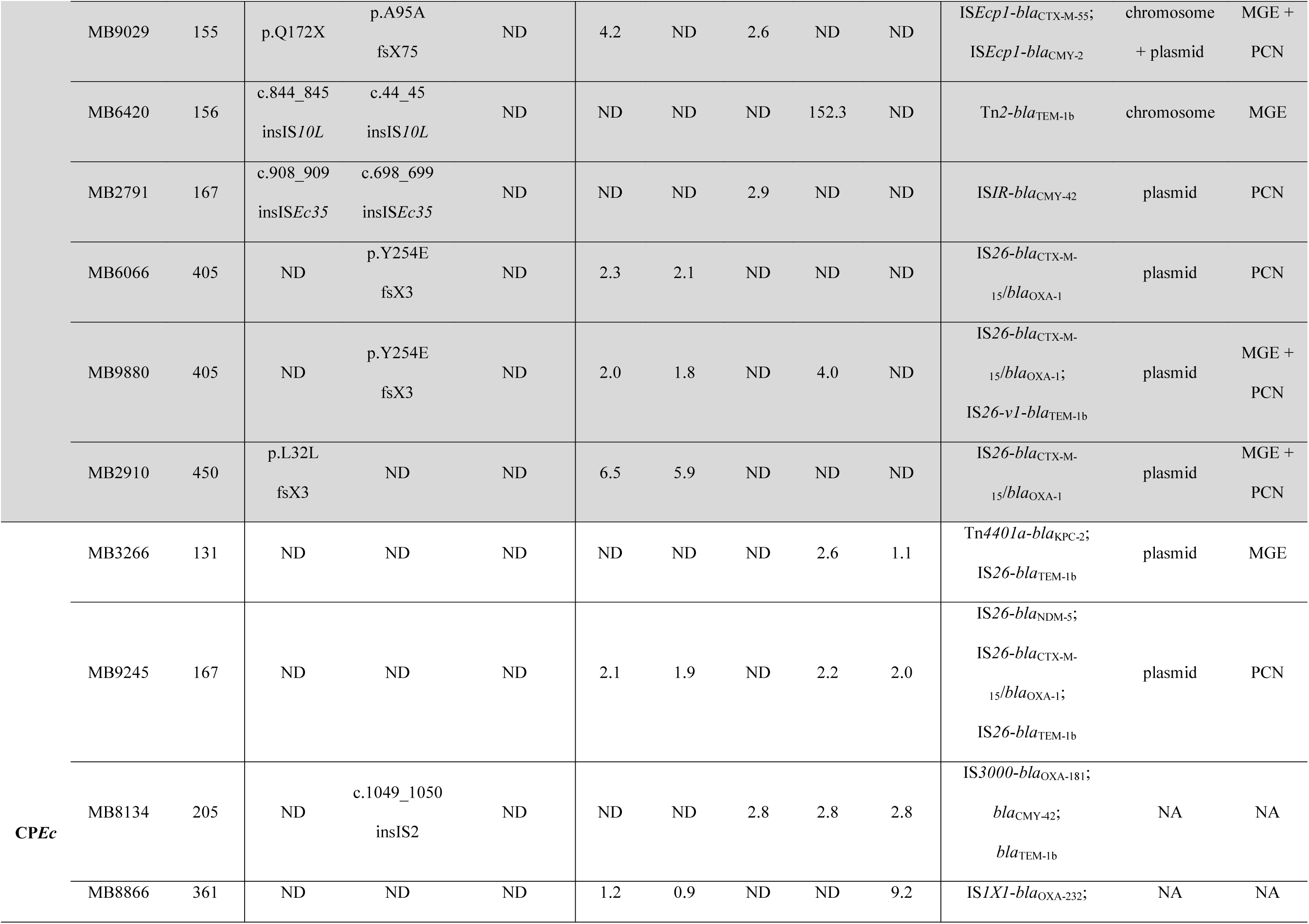
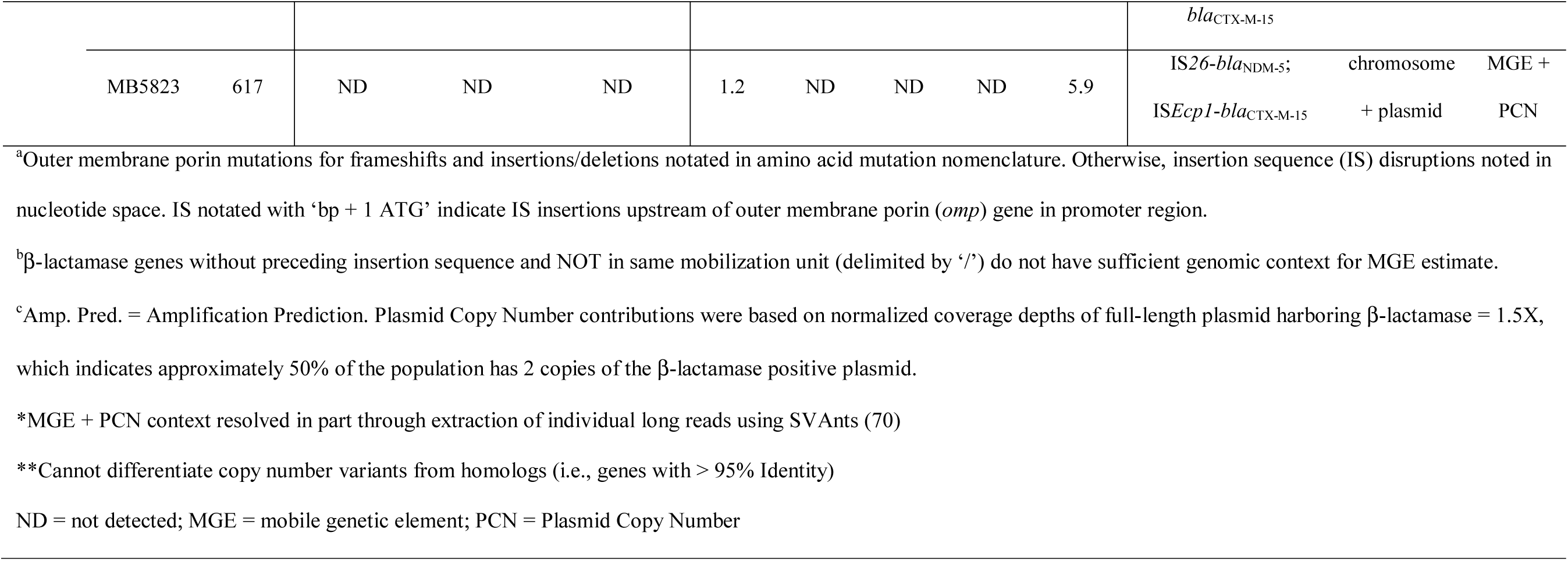
Summary of Carbapenem Non-Susceptibility Mechanisms for *E. coli* with β-Lactamase Amplifications.

**Table 2:**
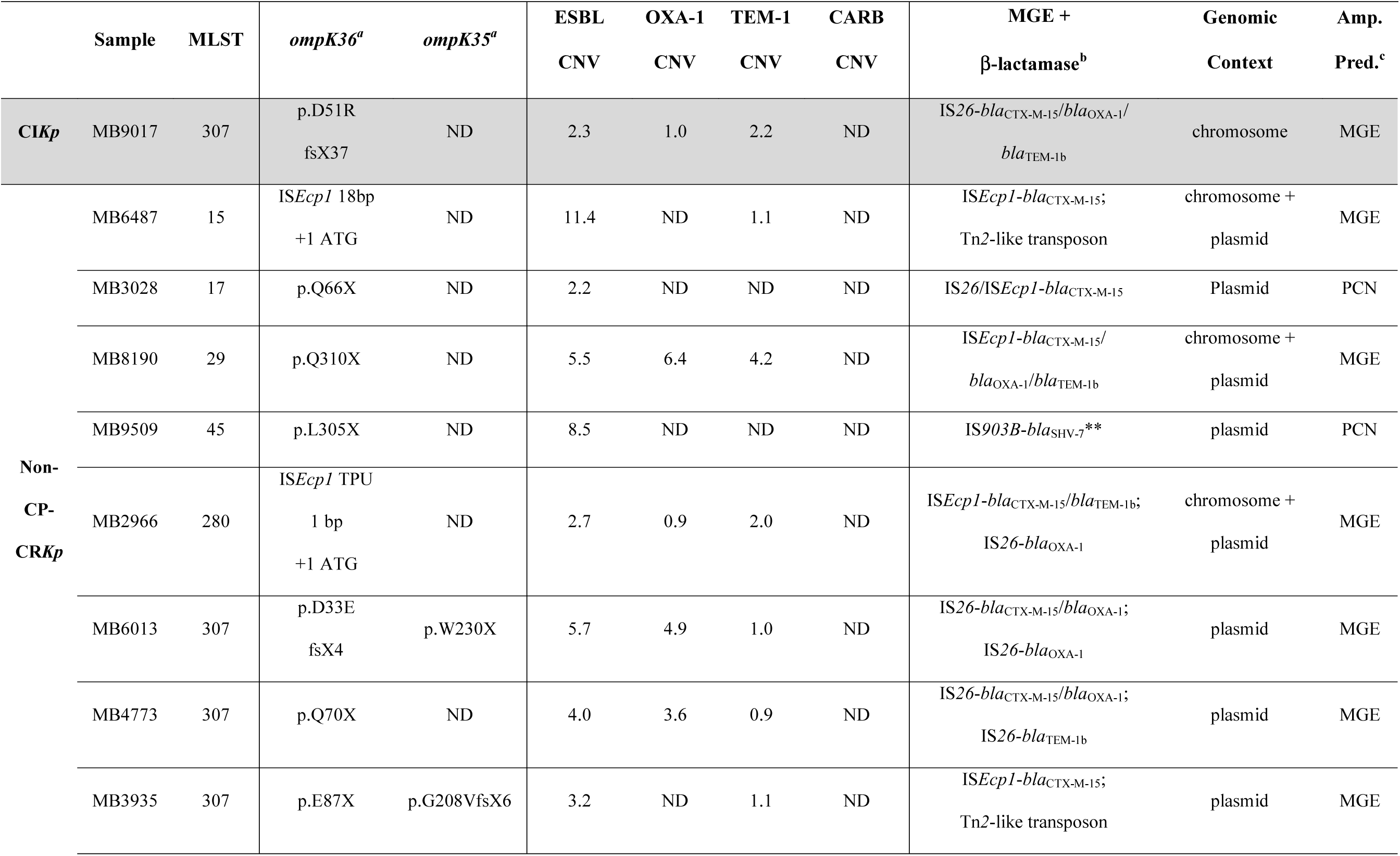
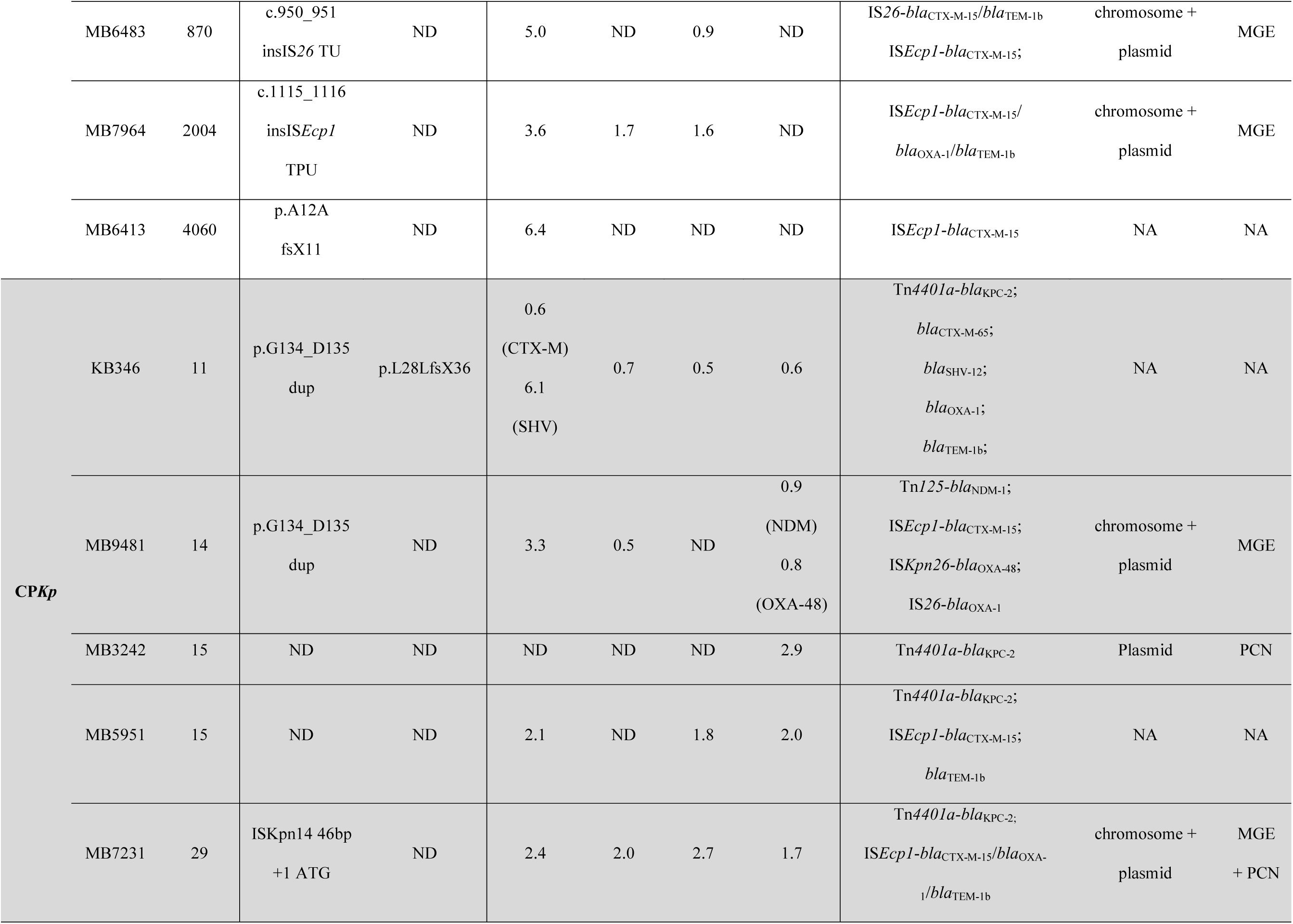
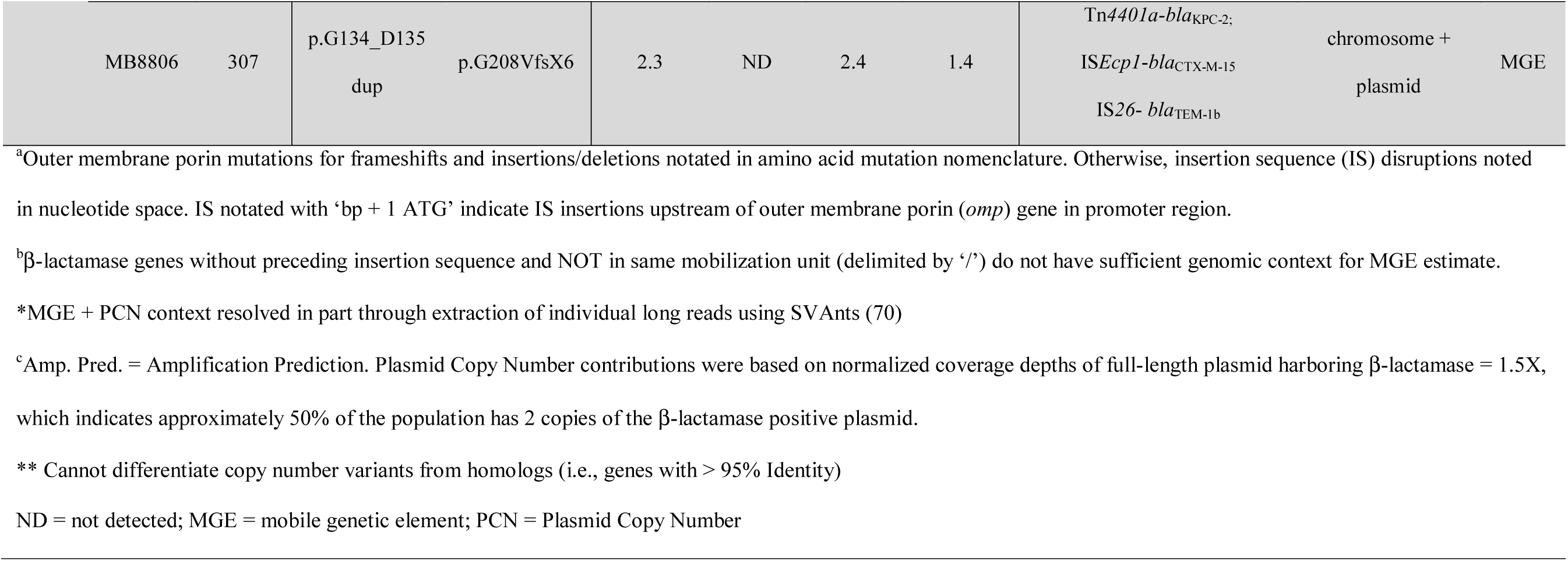
Summary of Carbapenem Non-Susceptibility Mechanisms for *K. pneumoniae* with β-Lactamase Amplifications.

When considering the most commonly observed β-lactamase amplifications, we often detected the syntenic coupling on MGEs of *bla*_OXA-1_ and/or *bla*_CTX-M-15_ with frequent gene amplification either through a TPU or TU structure in CNS*Ec* (11/27; 41%) or CNS*Kp* (8/15;53%) as presented in **Table 1** and **Table 2** respectively. Indeed, when measuring binary presence/absence of β-lactamase genes in the entire CNSE cohort, 41% (32/79) of CNSE had *bla*_CTX-M-15_/*bla*_OXA-1_ co-carriage with both chromosomal and/or plasmid contexts. Of the 31 CNSE isolates that had ONT data available and *bla*_CTX-M-15_/*bla*_OXA-1_ co-carriage (*N.B.*, one of the *bla*_CTX-M-15_/*bla*_OXA-1_ positive isolates only had a draft assembly), six isolates (three *E. coli* and three *K. pneumoniae*) had the two genes co-localized solely on the chromosome. The majority of *bla*_CTX-M-15_/*bla*_OXA-1_ co-localization was observed in a plasmid context (81%; 25/31) with all but one CNSE isolate (MB5646) having co-carriage on multi-replicon IncF type plasmids. Therefore, we calculated an estimate of pairwise average nucleotide identity (ANI) of all IncF type plasmids harboring *bla*_CTX-M-15_*/bla*_OXA-1_ **(Figure 4)** to determine the relatedness of these IncF plasmids and see if there was evidence of interclade and interspecies transmission. A full-length visualization of the multi-replicon F type plasmids can be found on Supplemental Figure 3.

**Figure 4.**
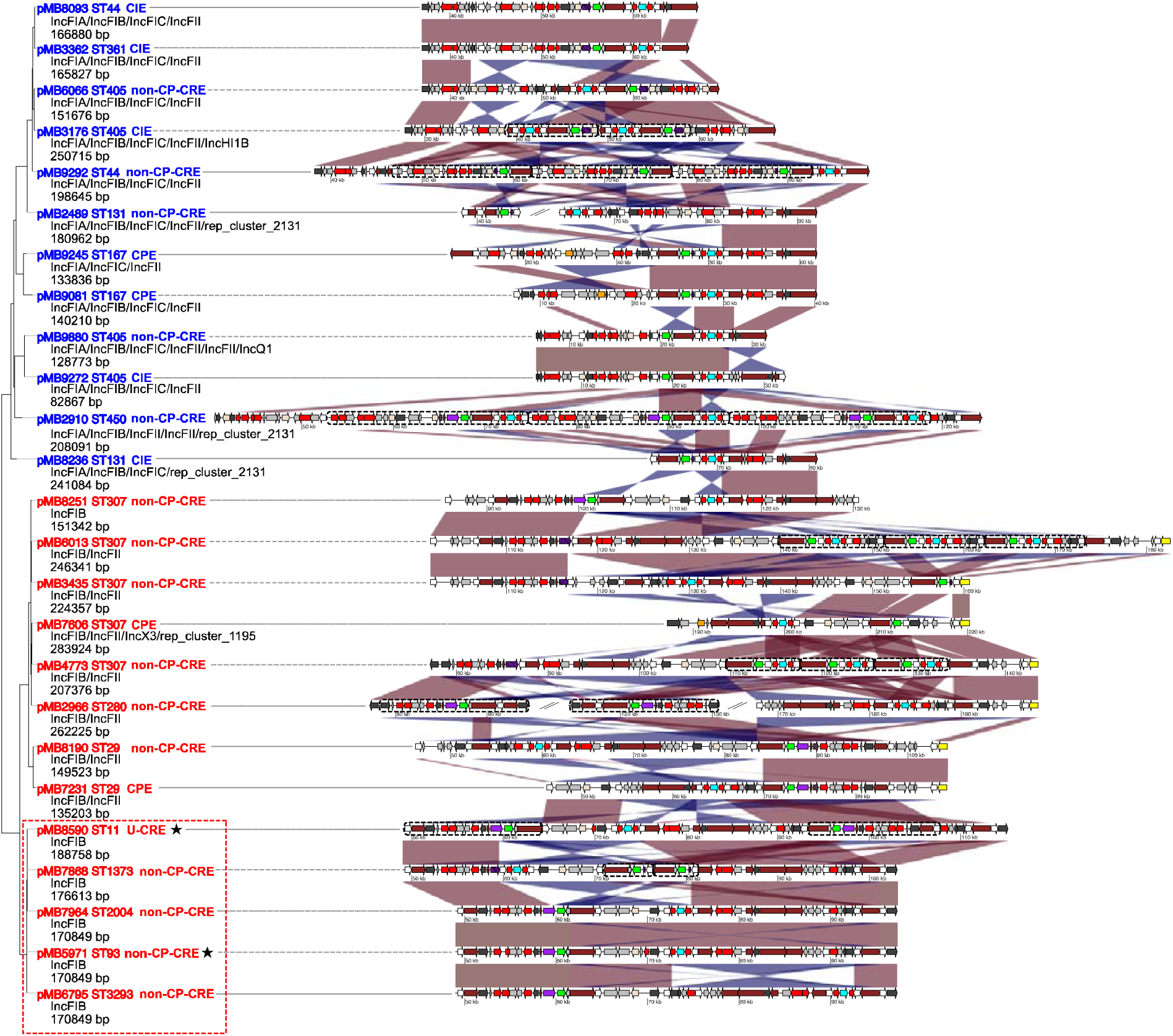
Multi-replicon IncF-type plasmids co-carrying *bla*_CTX-M-15_ and *bla*_OXA-1_ shared across multiple Enterobacterales species. Neighbor-joining (NJ) tree based on estimated ANI pairwise distances of full length, IncF-type multi-replicon plasmids with red tip labels indicating *Klebsiella* spp. and blue tip labels indicating *E. coli* plasmids. MOB Typing designations with plasmid size are beneath each respective NJ tree tip label. Regions of plasmid are subset from each respective plasmid with position indicated on each structure to highlight multidrug resistance region that includes *bla*_OXA-1_ (blue) and *bla*_CTX-M-15_ (green) open reading frame labels. Transposase/integrase (dark gray), IS*26* transposase (white), IS*26*-v1 (off-white), IS*Ecp1* transposase (purple), Tn3-like elements (brown), carbapenemases (orange), other antimicrobial genes (red), *rep* genes (yellow), and other genes (light gray) are labelled accordingly. Striped, purple IS*Ecp1* transposase ORFs indicate a disruption due to IS*26* or IS*26*-v1. Region on NJ tree enclosed by dotted red squares share ∼99% identity and with three plasmids (pMB7964, pMB5971, pMB6796) having ∼99% coverage. Stars adjacent to tip labels indicate non-*K. pneumoniae* species (pMB5971 = *K. aerogenes*; pMB8590 = *K. michiganensis*). Linear comparisons between sequences indicate homology shared (min length 1000 bp and >90% identity) in direct (red) and reverse (blue) orientation.

The ANI of all *bla*_CTX-M-15_*/bla*_OXA-1_ positive IncF type plasmids was highly similar (AVG = .94; SD = .04) across *E. coli* (n= 12) and *Klebsiella* spp. (n=13) with two primary clusters that formed by species when observing the Neighbor Joining distance inferred dendrogram **(Figure 4)**. The discrimination between *E. coli* and *K. pneumoniae* IncFIB plasmids was largely due to differences in transmission of well characterized replication initiation protein alleles found in *Klebsiella spp.* (*i.e*., IncFIB_K_) and *E. coli* (*i.e.*, IncFIB(AP001918)). One nested cluster of five IncFIB plasmids demarcated by a red box on **Figure 4** shared >99.9% ANI across three unique *K. pneumoniae* STs (pMB7868_1, pMB7964_1, pMB6795_1), *K. aerogenes* (pMB5971_1), and *K. michiganensis* (pMB8590_1). Interestingly, we observed *bla*_CTX-M-15_ and/or *bla*_OXA-1_ amplification occurring on 8/25 (32%) plasmids **(Figure 4; black striped boxes)** with all but one plasmid (pMB2966_1) having an IS*26* mediated TU amplification. Out of the 7 TUs with TU-mediated amplification, six were tandem arrays whereas only one plasmid (pMB8590_1) had a segmental duplication (i.e., mobilization to another genomic context) **(Figure 4)**.

We next sought to characterize and distinguish the IS*26*- and IS*Ecp1*-mediated mechanisms that were responsible for mobilizing *bla*_CTX-M-15_*/bla*_OXA-1_ from both a plasmid and chromosomal context. **Figure 5A** provides an illustration of a pseudo-compound transposon (PCT) that can be made of two or more IS*26* units, which must include flanking IS*26* transposase in direct orientation for potential co-integrate formation to occur and mobilize the passenger AMR genes (32). **Figure 5B** shows the highly modular, mosaic structures of these PCTs where except for one PCT (MB2910_PCT), these structures include an IS*26* or IS*26*-v1 element upstream of *bla*_CTX-M-15_, which inactivates IS*Ecp1* activity. Interestingly, these PCTs with inactivated IS*Ecp1* were more commonly observed in *E. coli* in contrast to *K. pneumoniae* apart from five isolates that all belonged to the emergent ST307 *K. pneumoniae* clade **(Figure 5B)**. There was also only one isolate (MB2489) with TU segmental duplication **(Figure 5B)** where based on chromosomal gene content present on the plasmid, there was likely a chromosome-to-plasmid IS*26* transposase-mediated cointegration formation event (33, 34).

**Figure 5.**
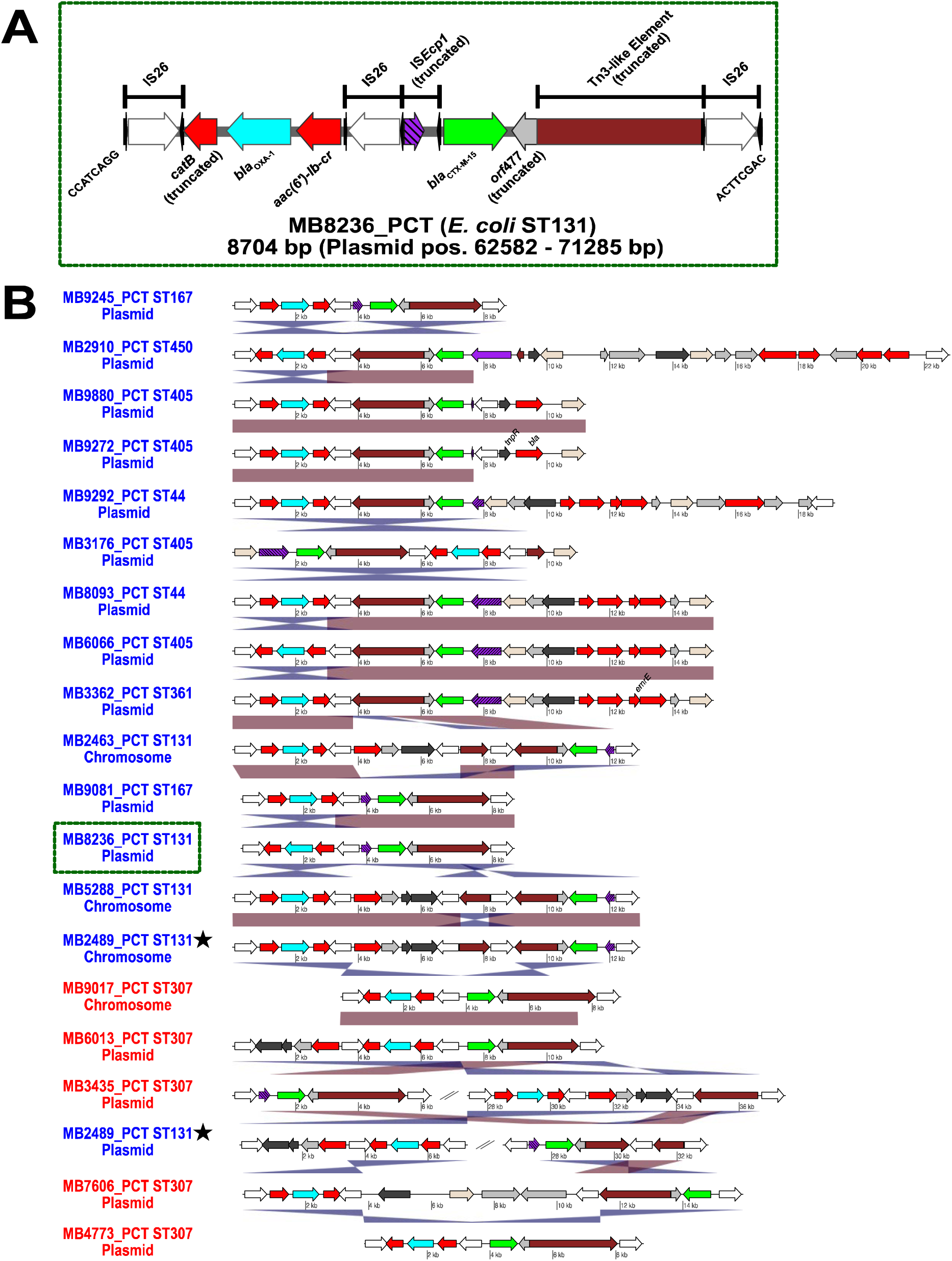
Pseudo-compound transposons (PCTs) driving mobilization and amplification of ESBL and narrow-spectrum. β**-lactamases.** Transposase/integrase (dark gray), IS*26* transposase (white), IS*26*-v1 (off-white), IS*Ecp1* transposase (purple), Tn3-like elements (brown), other antimicrobial genes (red), *bla*_OXA-1_ (blue), *bla*_CTX-M-15_ (green), and other genes (light gray) labelled accordingly. Striped, purple IS*Ecp1* transposase ORFs indicate a disruption due to IS*26* or IS*26*-v1. **(A)** Representation of pseudo-compound transposon (MB8236_PCT) flanked by IS*26* in direct orientation within a plasmid context. Black arrows flanking IS transposases indicate inverted repeats. 8-bp DNA flanking IS*26* on linearized representation of PCT. Position on plasmid indicated in parenthesis. **(B)** Plasmid and chromosomal contexts of PCT within *E. coli* (blue) and *K. pneumoniae* (red) indicating blastn identities as described in **Figure 4**. Stars indicate PCTs arising from the same genome. Green dotted line highlights the PCT that is fully annotated in **(A)**. Linear comparisons between sequences indicate homology shared (min length 1000 bp and >90% identity) in direct (red) and reverse (blue) orientation.

The other common MGE with the potential to mobilize *bla*_CTX-M-15_*/bla*_OXA-1_ was IS*Ecp1*-mediated transposable units (TPUs) with **Figure 6A** providing a schematic for a representative *K. pneumoniae* TPU (MB7231_TPU) found in a chromosomal context. In contrast to CNS*Ec*, 53% of FIB *Klebsiella* spp. plasmids had intact IS*Ecp1* immediately upstream of *bla*_CTX-M-15_ suggesting the potential for TPU formations as the primary driver of *bla*_CTX-M-15_ mobilization in non-ST307 CNS*Kp* **(Figure 6B)**. There were three CNS*Kp* isolates that had plasmid-to-chromosome transfer of IS*Ecp1* mediated TPUs as detected by 5bp target site duplications flanking the inverted repeat regions of the chromosomal TPUs **(Figure 6B)**. Taken together, our analysis highlights the enrichment of IS*26*/IS*Ecp1* structures present in CNSE that are overwhelmingly responsible for the amplifications of β-lactamase genes, in particular, *bla*_CTX-M-15_ and *bla*_OXA-1_ in our cohort.

**Figure 6.**
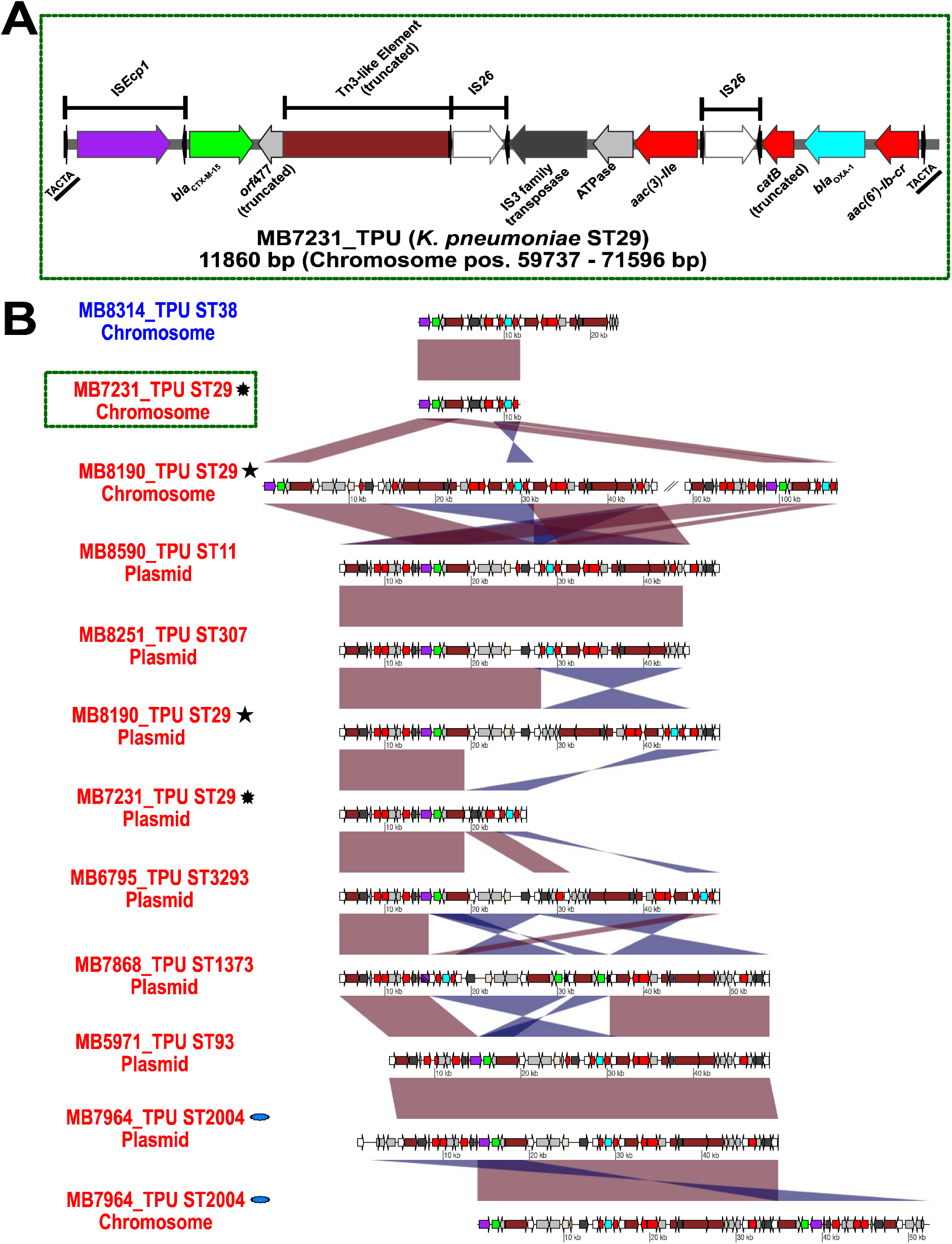
Transposition Units (TPUs) driving mobilization and amplification of ESBL and narrow-spectrum. β**-lactamases.** Transposase/integrase (dark gray), IS*26* transposase (white), IS*26*-v1 (off-white), IS*Ecp1* transposase (purple), Tn*3*-like elements (brown), other antimicrobial genes (red), *bla*_OXA-1_ (blue), *bla*_CTX-M-15_ (green), and other genes (light gray) labelled accordingly. Striped, purple IS*Ecp1* transposase ORFs indicate a disruption due to IS*26*. **(A)** Example of *K. pneumoniae* chromosomal context of transposition unit (MB7231_TPU) mobilized from plasmid-to-chromosome via IS*Ecp1.* Black arrows flanking IS transposases indicate inverted repeats. 5-bp direct repeat (underlined) flanking MB7231_TPU indicated on end of linearized representation of TPU. Position on chromosome indicated in parenthesis. **(B)** Plasmid and chromosomal contexts of TPU within *E. coli* (blue) and *K. pneumoniae* (red) indicating blastn identities as described in **Figure 4**. Matching symbols adjacent to labels indicate TPUs arising from the same genome. Green dotted line highlights the TPU that is fully annotated in **(A)**. Linear comparisons between sequences indicate homology shared (min length 1000 bp and >90% identity) in direct (red) and reverse (blue) orientation.

### Characterization of unconfirmed non-susceptible *Enterobacterales* (U-CNSE) isolates

In light of the increasing recognition of the impact of unconfirmed CRE (1), we next sought to characterize a subset of *E. coli* and *Klebsiella* spp. for which we had not confirmed carbapenem non-susceptibility to our non-CP-CNSE isolates. In marked contrast to non-CP-CR*Ec* and non-CP-CR*Kp* isolates, none of the U-CNS *E. coli* (n=3) and *Klebsiella* spp. (n=2) had mutated OmpC/OmpF (OmpK36/OmpK35) encoding genes **(Figure 2**; Table S2). All the U-CNS *E. coli* and *Klebsiella* spp. were *bla*_CTX-M_ positive (4 *bla*_CTX-M-15_; 1 *bla*_CTX-M-55_); furthermore, amplification of ESBL encoding enzymes were detected (median ESBL CNV = 2.4X) among all 5 U-CNS *E. coli* and *Klebsiella* spp. β-lactamase gene amplification in the U-CNS isolates shared similar mechanisms to that observed for the non-CP-CRE strains. For example, MB8590 (*K. michiganensis*; ST11) had evidence of a plasmid TU harboring *bla*_CTX-M-15_*/bla*_OXA-1_ that had two copies via segmental duplication **(Figure 4)**. Furthermore, this TU included an intact IS*Ecp1* **(Figure 6B)** suggesting the potential for TPU-mediated mobilization as well. Although a small number of isolates were examined, these data indicate that intact porins are the major distinction between unconfirmed and CNS *E. coli* and *Klebsiella* spp.

## DISCUSSION

Through a comprehensive, comparative genomics analysis on a diverse array of CNSE bacteremia isolates, we expanded the current understanding of the breadth of MGE-mediated mechanisms used to overcome carbapenems in clinically impactful *Enterobacterales* strains. By analyzing normalized coverage depths of β-lactamase encoding genes in conjunction with the detection of binary presence/absence of β-lactamases and *omp* genes, we show that amplification of ESBL genes as well as disruption of *omp* genes are commonly found amongst invasive non-CP-CRE. Additionally, our ONT long-read sequencing data allowed for full characterization of the complex MGE-mediated gene amplifications and genetic alterations that can generate carbapenem resistance in the absence of a carbapenemase. The increasing appreciation of both the scope and clinical impact of non-CP-CRE (1, 2) highlights the need to develop novel preventive and therapeutic strategies for this understudied group of organisms.

A key finding was the high prevalence of CNSE organisms that lacked carbapenemases with non-CP strains accounting for well over 70% of both CNS *E. coli* and *K. pneumoniae* in our cohort. One possible explanation for this finding was our inclusion of organisms with carbapenem intermediate susceptibility phenotypes (*i.e.,* CIE strains), a decision which was based on the recent CRACKLE-2 finding that patients with unconfirmed CNSE, which often tested intermediate to ertapenem or other carbapenems, had similar clinical outcomes to patients with confirmed CRE (1). Given that carbapenem MICs tend to be lower for non-CP-CRE vs. CPE (1, 28, 35), our inclusion of CIE strains likely increased our proportion of non-CP isolates. However, even when only CRE isolates were considered, we still observed a predominance of non-CP organisms for both *E. coli* (19/25, 76%) and *K. pneumoniae* (18/26, 69%). Whereas a high percentage of non-CP-CR*Ec* strains has consistently been found in CRE surveillance studies, the opposite is true of *K. pneumoniae* where the high prevalence of *bla*_KPC_ typically results in >70-80% of CR*Kp* organisms being carbapenemase positive in the United States (1, 28). The high percentage of non-CP-CRE in our cohort was particularly interesting given that we only examined bacteremia isolates which are sufficiently fit to cause a serious infection inasmuch as non-CP-CRE isolates are often considered to have a fitness defect relative to CPE strains (36–38). The reasons underlying the high prevalence of non-CP isolates in our bacteremia cohort are not currently known but may include relatively stringent infection control practices amongst our highly immunocompromised patients. Recently, Black et al. noted a higher prevalence of non-CP-CRE (59%) in south Texas where non-CP-CRE patients were more likely to receive a longer duration of antibiotic treatment as well as more likely to have an emergency department visit compared to CPE albeit with low number of observations (39). This finding is consistent with our cancer patient population that receives a high level of antibiotic treatment (40) and coincides with the finding that previous antibiotic exposure has been identified as a risk factor for non-CP-CRE relative to CPE in other studies (7).

The high percentage of non-CP organisms in our cohort led us to focus on using our genomic data to better understand mechanisms driving carbapenem resistance in the absence of a carbapenemase. There were several important findings from these analyses. First, consistent with previous data based primarily on laboratory studies of passaged strains and PCR based methods (22–24), we found that non-CP-CRE almost always had combined porin disruption and amplification of ESBL encoding genes. To our knowledge, our study is the first to systematically demonstrate ESBL gene copy number increase in a large cohort of non-CP-CRE bacteremia isolates. It is thought that the porin disruption limits carbapenem presence in the periplasm to the point where high level ESBL production can inactivate sufficient carbapenem to generate resistance (4). Thus, incorporating porin assessment and β-lactamase gene amplification could assist with predicting *Enterobacterales* carbapenem susceptibility using genomic data (41–43). Second, the non-CP-CRE isolates were genetically heterogenous and primarily encoded various CTX-M type ESBLs with or without OXA-1. ESBL variants of TEM or SHV were quite rare in *E. coli* (n=2) and *K. pneumoniae* (n = 3) as were plasmid-borne AmpC in *E. coli* (n=5) and not detected in *K. pneumoniae*. These findings may reflect the dominant nature of CTX-M containing strains amongst ESBL isolates and are congruent with a previous laboratory study showing that the type of CTX-M containing enzyme did not impact the rate of ertapenem resistance development under selective pressure (24). Finally, we observed minimal clonality amongst the non-CP-CRE strains indicating that the organisms developed carbapenem resistance independently rather than being transmitted between patients. This hypothesis is supported by our observation that in many of the non-CP-CR *E. coli* and *K. pneumoniae* cases, the patients had previously had a bloodstream infection with an ESBL producing, carbapenem-susceptible organism. Thus, it is highly likely that carbapenem treatment of the ESBL infection selected for non-CP-CRE strains via ESBL amplification and porin disruption. Given that in our previous study only a small percentage of patients treated for an ESBL infection subsequently developed a non-CP-CRE infection (13), we are actively investigating why particular genetic backgrounds may contribute to a higher probability of developing carbapenem resistance versus other ESBL positive *Enterobacterales* strains.

The use of ONT sequencing was critical in helping to delineate the diverse MGE mechanisms underlying increases in ESBL gene copy numbers, which in general are not discernable with the commonly used short-read, whole-genome sequencing or PCR-based approaches (15). The vast majority of the ESBL amplifications involved CTX-M encoding genes either mobilized by IS*26* ‘translocatable units’ or IS*Ecp1* ‘transposition units’ via segmental duplication or *in situ* ‘tandem’ amplification. Both IS*26* and IS*Ecp1* contain transposases capable of mobilizing AMR genes (albeit very different mechanisms), with IS*26* mediated gene amplification increasingly recognized as a cause of progressive resistance to various β-lactams (13, 17, 18, 33, 34, 44). The complex MGEs amplified by IS*26* and IS*Ecp1* often contained non-β-lactamase encoding genes that confer resistance to aminoglycosides (e.g., *aac(6’)-Ib-cr*), tetracyclines (e.g., *tetAR*), trimethroprim (e.g., *dfrA17*), and sulfonamides (e.g., *sul1*) as illustrated in **Figure 4 - 6.** Therefore, similar to CPE, our non-CP-CRE were often multi-drug resistant (Table S3), further hindering treatment options. Another finding of concern was identifying IS*26* or IS*Ecp1* co-amplification of two β-lactamases on the same transposable unit (**Table 1** and **Table 2)**, typically *bla*_CTX-M-15_ along with *bla*_OXA-1_ but also *bla*_CTX-M-15_ with *bla*_CMY-4_ and *bla*_CTX-M-55_ with *bla*_CMY-2_. These dual β-lactamase encoding gene amplified organisms often were non-susceptible to meropenem in addition to ertapenem (Table S3).

Our findings along with other data (43, 45, 46) suggest carbapenem non-susceptible *Enterobacterales* reside along a spectrum mediated to a major degree by changes in porin function and β-lactamase gene copy number. It is likely that unconfirmed CNSE consist of a heterogenous population of ESBL-positive, carbapenem-adapting strains with β-lactamase gene amplifications/porin disruptions which may give different phenotypic results depending on the particular colony tested (47). Further carbapenem adaptation may fix a single porin disruption as seen in our *E. coli* ST405 isolates in **Figure 2**, and/or increase in β-lactamase gene copy number within the population leading to a carbapenem intermediate phenotype that progresses to full resistance through further β-lactamase amplification and dual outer membrane porin disruption. This progressive β-lactam resistance model is analogous to that recently identified for *bla*_TEM-1_ and *bla*_OXA-1_ amplifications mediating piperacillin-tazobactam resistance (13, 16, 17, 48). The increasing rates of ESBL positive *Enterobacterales* infections means that there are growing opportunities for development of non-CP-CRE. Given the widespread nature of IS*26*-mediated TUs and IS*Ecp1-*mediated TPUs in association with ESBL enzymes, our data suggest that optimizing carbapenem therapy (choice of carbapenem, dose, and duration) of ESBL infections is likely to be critical to minimizing non-CP-CRE emergence.

Our study has some inherent limitations. First, we only assayed strains from a gDNA context. It is likely that non-CP-CRE mechanisms also include transcriptional and post-transcriptional changes that we did not discern. However, there were only a handful of CNSE strains where a DNA-based explanation for an observed phenotype could not be identified, and these strains will be assessed using other methodologies as part of future studies. Second, we focused on particular genomic areas, specifically known β-lactamase encoding elements and porin encoding genes. Thus, it remains possible that other, yet to be identified DNA alterations, contributed to the carbapenem susceptibility phenotypes. Similarly, we did not recreate the DNA modifications of interest in an isogenic background to conclusively demonstrate that the identified changes conferred carbapenem resistance. However, our findings are in line with those derived from previous laboratory passaged and genetically altered strains (22–24). Finally, given the large number of sequenced isolates, we did not assess for population heterogeneity, the impact of which we attempted to minimize by performing phenotypic and genotypic analyses on the same single colony.

In summary, we present a cohort of fully resolved genomes of carbapenem non-susceptible *Enterobacterales* causing invasive infections, focusing on a large number of non-carbapenemase producing *E. coli* and *K. pneumoniae* isolates. Our data shed light on the pleiotropic and potentially widespread mechanisms underlying the non-CP-CRE phenotype and suggest that antimicrobial stewardship practices are likely to be critical in efforts to decrease non-CP-CRE impact.

## MATERIALS AND METHODS

### Study Design

Our lab has a comprehensive storage of The University of Texas MD Anderson Cancer Center (MDACC) bacteremia isolates (*i.e.,* the Microbe Bank Database [MBD]) dating back to 2012 stocked at -80°C in thioglycolate media with 25% glycerol. CLSI 2018 M100 guidelines were used to determine MIC breakpoint interpretations for carbapenem resistance (49). *Enterobacterales* bacteremia isolates (n = 143) with a non-susceptible, MIC interpretation to ertapenem (ETP) (> 0.5 μg/mL) or meropenem (MEM) (> 1 μg/mL) as reported by the MDACC Division of Pathology and Laboratory Medicine (PLM) clinical microbiology laboratory were selected using the Epic® EHR software, workbench reporting tool from July 1^st^, 2016, to June 30^th^, 2020. *Enterobacterales* species with intrinsic resistance to carbapenems (*e.g., Proteus mirabilis*) were excluded from selection. Candidate isolates underwent additional MIC testing to confirm ETP non-susceptibility as identified by the PLM lab using ETest (bioMérieux) gradient MIC strips. Definitions of carbapenem non-susceptibility were based on the following criterion:

1. carbapenemase-producing *Enterobacterales* (CPE) = carbapenemase detection confirmed through whole genome sequencing (WGS);
2. non-carbapenemase-producing carbapenem resistant *Enterobacterales* (non-CP-CRE) = no carbapenemase detected in WGS with confirmation Etest ETP MIC ≥ 2 μg/mL AND MDACC ETP MIC ≥ 2 μg/mL OR MEM MIC ≥ 4 μg/mL;
3. carbapenem intermediate *Enterobacterales* (CIE) = (a) confirmation ETest 0.5 μg/mL < ETP MIC < 2.0 μg/mL OR (b) MDACC MIC where 0.5 μg/mL < ETP MIC < 2.0 μg/mL OR 1 μg/mL < MEM MIC < 4.0 μg/mL;
4. Unconfirmed carbapenem non-susceptible *Enterobacterales* (U-CNSE) = confirmation ETest ETP MIC≤0.5 μg/mL.

CNSE exclusion criteria included isolates not available in the MBD (n=10), serial isolates (*i.e.,* any consecutive, recurrent bacteremia isolate with identical species as identified by the PLM lab) (n=25), isolates from same culture (n=4), and U-CNSE phenotype isolates and/or isolates with no growth on ertapenem (0.5 μg/mL) supplemented THY agar (n=25). The first available ETP non-susceptible isolate per patient from the MBD that met the above definition, and the screening process was selected for whole genome sequencing. There were two isolates, MB8134 and MB8251, with differential *Enterobacterales* species cultured from the same patient and were isolated 18 days apart that were included in the total CNSE cohort. After screening for carbapenem non-susceptibility from available isolates (see Figure 1), our sampling frame resulted in 79 total CNSE isolates that were sequenced from 78 unique patients. In addition to our CNSE WGS cohort, we performed WGS on 8 U-CNSE to investigate unstable carbapenem non-susceptible phenotypes. An antibiogram of the 79 CNSE isolates + 8 U-CNSE isolates is available on Table S3.

### Illumina short-read and Oxford Nanopore Technologies long-read sequencing

All isolates were streaked from the MBD collection and grown on THY overnight at 37°C. Single colonies were picked and grown in LB broth for 4 hours at 37°C with mild agitation and subsequently a pellet was stored at -80°C until gDNA extraction. The extraction of gDNA was performed using the MasterPure™ Complete DNA and RNA purification Kit using manufacturer’s instructions. Genomic DNA concentration was measured using the Qubit™ 4 fluorometer with complementary measurement of concentration and A260/280; A260/230 performed on an Eppendorf BioPhotometer. Isolates were then library prepped using the Illumina DNA Prep kit and sequenced using the Illumina NovaSeq 6000 platform. Select isolates were then sequenced using the long-read Oxford Nanopore Technologies (ONT) GridION platform with the Rapid Sequencing Kit (SQK-RAD004) per manufacturer’s instructions.

Short-read Illumina fastq data was trimmed, QC, and assembled using a customized workflow (Shropshire W, SPAdes_pipeline-v0.1.0-alpha, GitHub: https://github.com/wshropshire/SPAdes_pipeline) with assemblies generated using SPAdes-v3.15.3 using the ‘—isolate’ parameter in addition to default parameters for paired-end short-read data. Short-read and long-read were used with the Flye (Flye-2.9-b1768) assembler pipeline (Shropshire W, flye_hybrid_assembly_pipeline-v0.3.0-alpha, GitHub: https://github.com/wshropshire/flye_hybrid_assembly_pipeline). Genome assembly quality was assessed with CheckM-v1.2.0 (50) with mean coverage depth of complete and draft assemblies calculated using mosdepth-v0.3.3 (51). An overview of genome assembly quality metrics is presented on Table S5.

### Pan genome and maximum likelihood (ML) phylogenetic analysis

Complete and draft assemblies were then used as input for pan genome analysis using Panaroo-v.1.2.9 (52) using the ‘moderate’ –clean-mode parameter with the mafft core gene alignment option. This core gene alignment file was then used as input to create a maximum-likelihood phylogenetic tree with IQTree2-2.2.0-beta (53). When creating the core gene inferred ML phylogenetic tree, model selection was performed using ModelFinder (54), a non-parametric bootstrap approximation, UFBoot (55) (n=1000), and an SH-aLRT (n=1000) test to further evaluate branch lengths. Tree visualization along with the addition of metadata was completed using ggree-3.1.1 and ggtreeExtra-1.0.4 respectively. Clustering of isolates based on core gene alignment was assessed using the rhierhaps-1.1.3 tool (56). Pairwise SNP differences were assessed using the snp-dists (Seemann T, snp-dists-v0.8.2 https://github.com/tseemann/snp-dists).

### Antimicrobial resistance gene and *in silico* typing profiles

Kleborate-v2.0.4 (57) was used with draft and complete assemblies to identify K and O antigen profiles (Kleborate confidence scores of ‘Good’ or better), MLST, acquired and chromosomal antimicrobial resistance, and virulence factors for isolates belonging to the *Klebsiella pneumoniae* species complex (KpSC). Additionally, Kleborate (57) was used to designate species taxa for all isolates sequenced by calculating pairwise Mash distances (58) between each respective genome assembly and their Enterobacterales reference genomes (n=2619). All isolates had strong species matches (*i.e.,* Mash distances < 0.02). SerotypeFinder-2.0 Server (59) was used for *in silico* serotyping of *E. coli* isolates using a 85% ID/60% minimum length threshold for O and H antigen identification. Novel MLST schema not identified using Kleborate-v2.0.4 or the mlst-2.19.0 Perl script (Seemann T, mlst-2.19.0, GitHub: https://github.com/tseemann/mlst) was identified using the MLST-2.0 Server (60). Phylogroups of *E. coli* were detected using the ClermonTyping-20.03 tool (61) using the clermonTyping.sh script. The BLASTn alignment tool (BLAST 2.11.0+) was used with an in-house database of *E. coli ompC* and *ompF* genes (MG1655 K-12 reference) and their respective enterobacterial homologs identified in *Klebsiella* spp., *Enterobacter* spp., *Citrobacter* spp., and *Serratia marcescens* to characterize potential osmoporin gene disruption. SnapGene v5.0.8 was used to visualize these osmoporin gene disruptions and further characterize MGE associated insertions within the open reading frame and/or promoter region using ISFinder (62).

### AMR gene and plasmid copy number variation estimation

Antimicrobial resistance genes were detected using the KmerResistance-2.2.0 (63, 64) tool which uses KMA-1.3.24a to detect AMR genes using a short-read k-mer based alignment against the ResFinder (Accessed November 5^th^, 2021). These ResFinder hits were then used as input for a copy number variant estimation tool (Shropshire W, convict-v1.0, GitHub: https://github.com/wshropshire/convict), which estimates gene copy number variants by normalizing coverage depths to housekeeping genes. Genes present in >99% of the consensus, pan genome fasta file generated from Panaroo were used to control coverage depth. SVAnts (Hanson, B. GitHub: https://github.com/EpiBlake/SVants) was used to confirm copy number variants with individual ONT long-reads containing multiple tandem repeats of IS*26* and IS*Ecp1* multi-resistance determinant regions for isolates with increased coverage depth mapping visualized in IGV-2.9.4. A ratio of mean coverage depths of plasmid-to-chromosome was calculated using bwa mem alignments and the pileup.sh script from bbmap-v38.79 to get an approximation of plasmid copy number (PCN).

Plasmid typing of completed assemblies was completed using the mob_typer-v3.0.0 command line tool (65). FastANI-v1.31 (66) was used to estimate average nucleotide identity across plasmid and MGE structures with default settings. The bacsort script (Wick, R. GitHub: https://github.com/rrwick/Bacsort), ‘pairwise_identities_to_distance_matrix.py’ is used to convert FastANI pairwise distances to a distance matrix in PHYLIP format with a maximum genetic distance of 0.20. This distance matrix was used as input to create a neighbor-joining tree using the BIONJ algorithm (67) using the ape-v.5.6-1 R package (68). Genome comparisons and annotations of plasmid and MGE structures was performed using the genoPlotR-v0.8.11 R package (69). In order to filter multiple IS comparisons, a minimum sequence fragment length of 1000 bp was used to compare blastn identities ≥90% in direct (red) or reverse (blue) orientation.

### Statistics

All statistics were performed using R version 4.0.4 (2021-02-15). Significant increases in AMR gene copy numbers were assessed using one-sample Wilcoxon Tests with a one-sided, alternative hypothesis that mean CNV was greater than 1. Scatterplot and boxplots generated using ggplot2-3.3.5.

### Data Availability

Short-read Illumina data, long-read ONT data, complete and draft assemblies are available in the NCBI BioProject PRJNA836696 repository. Three samples (MB2315, MB2446, MB2463) have data available from a previous BioProject (PRJNA603908).

## ACKNOWLEDGEMENTS

Illumina short-read sequencing was done through the MDACC Advanced Technology Core (ATGC) using core grant CA016672(ATGC) with the Illumina NovaSeq6000 (NIH 1S10OD024977-01). Support for this study was provided by the National Institute of Allergy and Infectious Diseases (NIAID) R21AI151536 and P01AI152999 for S.A.S. NIAID K24AI121296, R01AI134637, R01AI148342, R21AI143229, a UTHealth Presidential Award, University of Texas System STARS Award, and Texas Medical Center Health Policy Institute Funding Program supported C.A.A. B.M.H. was supported by the NIAID K01AI148593-01. The research in the A.K. laboratory is supported by NIGMS 1R01GM133904-01 and the Welch Foundation Research Grant AU-1998-20190330.

## REFERENCES

1. van Duin D, Arias CA, Komarow L, Chen L, Hanson BM, Weston G, Cober E, Garner OB, Jacob JT, Satlin MJ, Fries BC, Garcia-Diaz J, Doi Y, Dhar S, Kaye KS, Earley M, Hujer AM, Hujer KM, Domitrovic TN, Shropshire WC, Dinh A, Manca C, Luterbach CL, Wang M, Paterson DL, Banerjee R, Patel R, Evans S, Hill C, Arias R, Chambers HF, Fowler VG, Kreiswirth BN, Bonomo RA. 2020. Molecular and clinical epidemiology of carbapenem-resistant Enterobacterales in the USA (CRACKLE-2): a prospective cohort study. The Lancet Infectious Diseases 20:731–741.

2. Guh AY, Bulens SN, Mu Y, Jacob JT, Reno J, Scott J, Wilson LE, Vaeth E, Lynfield R, Shaw KM, Vagnone PM, Bamberg WM, Janelle SJ, Dumyati G, Concannon C, Beldavs Z, Cunningham M, Cassidy PM, Phipps EC, Kenslow N, Travis T, Lonsway D, Rasheed JK, Limbago BM, Kallen AJ. 2015. Epidemiology of Carbapenem-Resistant Enterobacteriaceae in 7 US Communities, 2012-2013. JAMA 314:1479–87.

3. Dagher C, Salloum T, Alousi S, Arabaghian H, Araj GF, Tokajian S. 2018. Molecular characterization of Carbapenem resistant Escherichia coli recovered from a tertiary hospital in Lebanon. PLOS ONE 13:e0203323.

4. Nordmann P, Poirel L. 2019. Epidemiology and Diagnostics of Carbapenem Resistance in Gram-negative Bacteria. Clinical Infectious Diseases 69:S521–S528.

5. Satlin MJ, Jenkins SG, Walsh TJ. 2014. The global challenge of carbapenem-resistant Enterobacteriaceae in transplant recipients and patients with hematologic malignancies. Clin Infect Dis 58:1274–83.

6. Nguyen MH, Shields RK, Chen L, Pasculle AW, Hao B, Cheng S, Sun J, Kline EG, Kreiswirth BN, Clancy CJ. 2021. Molecular epidemiology, natural history and long-term outcomes of multi-drug resistant Enterobacterales colonization and infections among solid organ transplant recipients. Clinical Infectious Diseases doi:10.1093/cid/ciab427.

7. Marimuthu K, Ng OT, Cherng BPZ, Fong RKC, Pada SK, De PP, Ooi ST, Smitasin N, Thoon KC, Krishnan PU, Ang MLT, Chan DSG, Kwa ALH, Deepak RN, Chan YK, Chan YFZ, Huan X, Zaw Linn K, Tee NWS, Tan TY, Koh TH, Lin RTP, Hsu LY, Sengupta S, Paterson DL, Perencevich E, Harbarth S, Teo J, Venkatachalam I, Cherng B, Su Gin DC, Rama Narayana D, De PP, Li Yang H, Venkatachalam I, Teo J, Marimuthu K, Tse Hsien K, Tee N, Smitasin N, Oon Tek N, Say Tat O, Krishnan PU, Fong R, Lin Tzer Pin R, Pada SK, Thean Yen T, Koh Cheng T. 2019. Antecedent Carbapenem Exposure as a Risk Factor for Non-Carbapenemase-Producing Carbapenem-Resistant Enterobacteriaceae and Carbapenemase-Producing Enterobacteriaceae. Antimicrobial Agents and Chemotherapy 63.

8. Bouganim R, Dykman L, Fakeh O, Motro Y, Oren R, Daniel C, Lazarovitch T, Zaidenstein R, Moran-Gilad J, Marchaim D. 2020. The Clinical and Molecular Epidemiology of Noncarbapenemase-Producing Carbapenem-Resistant Enterobacteriaceae: A Case-Case-Control Matched Analysis. Open Forum Infect Dis 7:ofaa299.

9. Shropshire WC, Dinh AQ, Earley M, Komarow L, Panesso D, Rydell K, Gómez-Villegas SI, Miao H, Hill C, Chen L. 2021. Accessory Genomes Drive Independent Spread of Carbapenem-Resistant Klebsiella pneumoniae Clonal Groups 258 and 307. medRxiv.

10. Wyres KL, Lam MMC, Holt KE. 2020. Population genomics of Klebsiella pneumoniae. Nat Rev Microbiol 18:344–359.

11. Mathers AJ, Peirano G, Pitout JDD. 2015. The Role of Epidemic Resistance Plasmids and International High-Risk Clones in the Spread of Multidrug-Resistant Enterobacteriaceae. Clinical Microbiology Reviews 28:565–591.

12. Poirel L, Bonnin RA, Nordmann P. 2012. Genetic support and diversity of acquired extended-spectrum β-lactamases in Gram-negative rods. Infection, Genetics and Evolution 12:883–893.

13. Shropshire WC, Aitken SL, Pifer R, Kim J, Bhatti MM, Li X, Kalia A, Galloway-Peña J, Sahasrabhojane P, Arias CA, Greenberg DE, Hanson BM, Shelburne SA. 2020. IS26-mediated amplification of blaOXA-1 and blaCTX-M-15 with concurrent outer membrane porin disruption associated with de novo carbapenem resistance in a recurrent bacteraemia cohort. Journal of Antimicrobial Chemotherapy doi:10.1093/jac/dkaa447.

14. Khan A, Shropshire WC, Hanson B, Dinh AQ, Wanger A, Ostrosky-Zeichner L, Arias CA, Miller WR. 2019. Simultaneous Infection with Enterobacteriaceae and Pseudomonas aeruginosa Harboring Multiple Carbapenemases in a Returning Traveler Colonized with Candida auris. Antimicrobial Agents and Chemotherapy 64.

15. George S, Pankhurst L, Hubbard A, Votintseva A, Stoesser N, Sheppard AE, Mathers A, Norris R, Navickaite I, Eaton C, Iqbal Z, Crook DW, Phan HTT. 2017. Resolving plasmid structures in Enterobacteriaceae using the MinION nanopore sequencer: assessment of MinION and MinION/Illumina hybrid data assembly approaches. Microb Genom 3:e000118.

16. Schechter LM, Creely DP, Garner CD, Shortridge D, Nguyen H, Chen L, Hanson BM, Sodergren E, Weinstock GM, Dunne WM. 2018. Extensive gene amplification as a mechanism for piperacillin-tazobactam resistance in Escherichia coli. MBio 9:e00583–18.

17. Hubbard ATM, Mason J, Roberts P, Parry CM, Corless C, Van Aartsen J, Howard A, Bulgasim I, Fraser AJ, Adams ER, Roberts AP, Edwards T. 2020. Piperacillin/tazobactam resistance in a clinical isolate of Escherichia coli due to IS26-mediated amplification of blaTEM-1B. Nature Communications 11.

18. Feng Y, Liu L, McNally A, Zong Z. 2018. Coexistence of Two blaNDM-5 Genes on an IncF Plasmid as Revealed by Nanopore Sequencing. Antimicrob Agents Chemother 62.

19. Goodman KES, P J; Tamma, P D; Milstone, A M. 2016. Infection control implications of heterogeneous resistance mechanisms in carbapenem-resistant Enterobacteriaceae (CRE). Expert Rev Anti Infect Ther 14:95–108.

20. Mathers A. 2016. Mobilization of Carbapenemase-Mediated Resistance in Enterobacteriaceae. Microbiol Spectr 4.

21. Cuzon G, Naas T, Nordmann P. 2011. Functional characterization of Tn4401, a Tn3-based transposon involved in blaKPC gene mobilization. Antimicrob Agents Chemother 55:5370–3.

22. Tangden T, Adler M, Cars O, Sandegren L, Lowdin E. 2013. Frequent emergence of porin-deficient subpopulations with reduced carbapenem susceptibility in ESBL-producing Escherichia coli during exposure to ertapenem in an in vitro pharmacokinetic model. J Antimicrob Chemother 68:1319–26.

23. Adler M, Anjum M, Andersson DI, Sandegren L. 2013. Influence of acquired beta-lactamases on the evolution of spontaneous carbapenem resistance in Escherichia coli. J Antimicrob Chemother 68:51–9.

24. Girlich D, Poirel L, Nordmann P. 2009. CTX-M expression and selection of ertapenem resistance in Klebsiella pneumoniae and Escherichia coli. Antimicrob Agents Chemother 53:832–4.

25. Goessens WH, van der Bij AK, van Boxtel R, Pitout JD, van Ulsen P, Melles DC, Tommassen J. 2013. Antibiotic trapping by plasmid-encoded CMY-2 beta-lactamase combined with reduced outer membrane permeability as a mechanism of carbapenem resistance in Escherichia coli. Antimicrob Agents Chemother 57:3941–9.

26. van Boxtel R, Wattel, A. A., Arenas, J., Goessens, W. H., & Tommassen, J. 2017. Acquisition of Carbapenem Resistance by Plasmid-Encoded-AmpC-Expressing Escherichia coli. Antimicrobial agents and chemotherapy 61:e01413–16.

27. Partridge SR, Kwong, S. M., Firth, N., S.O. 2018. Mobile Genetic Elements Associated with Antimicrobial Resistance. Clin Microbiol Rev 31:e00088–17.

28. Karlsson M, Lutgring JD, Ansari U, Lawsin A, Albrecht V, McAllister G, Daniels J, Lonsway D, McKay S, Beldavs Z, Bower C, Dumyati G, Gross A, Jacob J, Janelle S, Kainer MA, Lynfield R, Phipps EC, Schutz K, Wilson L, Witwer ML, Bulens SN, Walters MS, Duffy N, Kallen AJ, Elkins CA, Rasheed JK. 2022. Molecular Characterization of Carbapenem-Resistant Enterobacterales Collected in the United States. Microb Drug Resist doi:10.1089/mdr.2021.0106.

29. Guérin F, Isnard C, Cattoir V, Giard JC. 2015. Complex Regulation Pathways of AmpC-Mediated β-Lactam Resistance in Enterobacter cloacae Complex. Antimicrobial Agents and Chemotherapy 59:7753–7761.

30. Clermont O, Christenson JK, Denamur E, Gordon DM. 2013. The Clermont Escherichia coli phylo-typing method revisited: improvement of specificity and detection of new phylo-groups. Environmental Microbiology Reports 5:58–65.

31. Hall MN, Silhavy TJ. 1981. The ompB locus and the regulation of the major outer membrane porin proteins of Escherichia coli K12. Journal of Molecular Biology 146:23–43.

32. Harmer CJ, Pong CH, Hall RM. 2020. Structures bounded by directly-oriented members of the IS26 family are pseudo-compound transposons. Plasmid 111:102530.

33. Harmer CJ, Moran RA, Hall RM. 2014. Movement of IS26-associated antibiotic resistance genes occurs via a translocatable unit that includes a single IS26 and preferentially inserts adjacent to another IS26. MBio 5:e01801–14.

34. Harmer CJ, Hall RM. 2016. IS26-Mediated Formation of Transposons Carrying Antibiotic Resistance Genes. mSphere 1.

35. David S, Reuter S, Harris SR, Glasner C, Feltwell T, Argimon S, Abudahab K, Goater R, Giani T, Errico G, Aspbury M, Sjunnebo S, Eu SWG, Group ES, Feil EJ, Rossolini GM, Aanensen DM, Grundmann H. 2019. Epidemic of carbapenem-resistant Klebsiella pneumoniae in Europe is driven by nosocomial spread. Nat Microbiol doi:10.1038/s41564-019-0492-8.

36. Buckner MM, Saw HT, Osagie RN, McNally A, Ricci V, Wand ME, Woodford N, Ivens A, Webber MA, Piddock LJ. 2018. Clinically relevant plasmid-host interactions indicate that transcriptional and not genomic modifications ameliorate fitness costs of Klebsiella pneumoniae carbapenemase-carrying plasmids. MBio 9:e02303–17.

37. Knopp M, Andersson DI. 2015. Amelioration of the Fitness Costs of Antibiotic Resistance Due To Reduced Outer Membrane Permeability by Upregulation of Alternative Porins. Mol Biol Evol 32:3252–63.

38. Adler M, Anjum M, Berg OG, Andersson DI, Sandegren L. 2014. High fitness costs and instability of gene duplications reduce rates of evolution of new genes by duplication-divergence mechanisms. Mol Biol Evol 31:1526–35.

39. Black CA, So W, Dallas SS, Gawrys G, Benavides R, Aguilar S, Chen C-J, Shurko JF, Lee GC. 2021. Predominance of Non-carbapenemase Producing Carbapenem-Resistant Enterobacterales in South Texas. Frontiers in Microbiology 11:3629.

40. Webb BJ, Majers J, Healy R, Jones PB, Butler AM, Snow G, Forsyth S, Lopansri BK, Ford CD, Hoda D. 2020. Antimicrobial Stewardship in a Hematological Malignancy Unit: Carbapenem Reduction and Decreased Vancomycin-Resistant Enterococcus Infection. Clinical Infectious Diseases 71:960–967.

41. Avershina E, Sharma P, Taxt AM, Singh H, Frye SA, Paul K, Kapil A, Naseer U, Kaur P, Ahmad R. 2021. AMR-Diag: Neural network based genotype-to-phenotype prediction of resistance towards beta-lactams in Escherichia coli and Klebsiella pneumoniae. Comput Struct Biotechnol J 19:1896–1906.

42. Tamma PD, Fan Y, Bergman Y, Pertea G, Kazmi AQ, Lewis S, Carroll KC, Schatz MC, Timp W, Simner PJ. 2019. Applying Rapid Whole-Genome Sequencing To Predict Phenotypic Antimicrobial Susceptibility Testing Results among Carbapenem-Resistant Klebsiella pneumoniae Clinical Isolates. Antimicrobial Agents and Chemotherapy 63.

43. Bulman ZP, Krapp F, Pincus NB, Wenzler E, Murphy KR, Qi C, Ozer EA, Hauser AR. 2021. Genomic Features Associated with the Degree of Phenotypic Resistance to Carbapenems in Carbapenem-Resistant Klebsiella pneumoniae. mSystems 6.

44. Zienkiewicz M, Kern-Zdanowicz I, Carattoli A, Gniadkowski M, Ceglowski P. 2013. Tandem multiplication of the IS26-flanked amplicon with the bla(SHV-5) gene within plasmid p1658/97. FEMS Microbiol Lett 341:27–36.

45. Patiño-Navarrete R, Rosinski-Chupin I, Cabanel N, Gauthier L, Takissian J, Madec J-Y, Hamze M, Bonnin RA, Naas T, Glaser P. 2020. Stepwise evolution and convergent recombination underlie the global dissemination of carbapenemase-producing Escherichia coli. Genome Medicine 12.

46. Ma P, He LL, Pironti A, Laibinis HH, Ernst CM, Manson AL, Bhattacharyya RP, Earl AM, Livny J, Hung DT. 2021. Genetic determinants facilitating the evolution of resistance to carbapenem antibiotics. eLife 10.

47. Nicoloff H, Hjort K, Levin BR, Andersson DI. 2019. The high prevalence of antibiotic heteroresistance in pathogenic bacteria is mainly caused by gene amplification. Nat Microbiol 4:504–514.

48. Rodriguez-Villodres A, Gil-Marques ML, Alvarez-Marin R, Bonnin RA, Pachon-Ibanez ME, Aguilar-Guisado M, Naas T, Aznar J, Pachon J, Lepe JA, Smani Y. 2019. Extended-spectrum resistance to beta-lactams/beta-lactamase inhibitors (ESRI) evolved from low-level resistant Escherichia coli. J Antimicrob Chemother doi:10.1093/jac/dkz393.

49. CLSI. 2018. Performance Standards for Antimicrobial Susceptibility Testing. Clinical and Laboratory Standards Institute, Wayne, PA.

50. Parks DH, Imelfort M, Skennerton CT, Hugenholtz P, Tyson GW. 2015. CheckM: assessing the quality of microbial genomes recovered from isolates, single cells, and metagenomes. Genome research 25:1043–1055.

51. Pedersen BS, Quinlan AR. 2018. Mosdepth: quick coverage calculation for genomes and exomes. Bioinformatics 34:867–868.

52. Tonkin-Hill G, Macalasdair N, Ruis C, Weimann A, Horesh G, Lees JA, Gladstone RA, Lo S, Beaudoin C, Floto RA, Frost SDW, Corander J, Bentley SD, Parkhill J. 2020. Producing polished prokaryotic pangenomes with the Panaroo pipeline. Genome Biology 21.

53. Minh BQ, Schmidt HA, Chernomor O, Schrempf D, Woodhams MD, von Haeseler A, Lanfear R. 2020. IQ-TREE 2: New Models and Efficient Methods for Phylogenetic Inference in the Genomic Era. Mol Biol Evol 37:1530–1534.

54. Kalyaanamoorthy S, Minh BQ, Wong TKF, Von Haeseler A, Jermiin LS. 2017. ModelFinder: fast model selection for accurate phylogenetic estimates. Nature Methods 14:587–589.

55. Hoang DT, Chernomor O, Von Haeseler A, Minh BQ, Vinh LS. 2018. UFBoot2: Improving the Ultrafast Bootstrap Approximation. Molecular Biology and Evolution 35:518–522.

56. Tonkin-Hill G, Lees JA, Bentley SD, Frost SDW, Corander J. 2018. RhierBAPS: An R implementation of the population clustering algorithm hierBAPS. Wellcome Open Res 3:93.

57. Lam MMC, Wick RR, Watts SC, Cerdeira LT, Wyres KL, Holt KE. 2021. A genomic surveillance framework and genotyping tool for Klebsiella pneumoniae and its related species complex. Nature Communications 12.

58. Ondov BD, Treangen TJ, Melsted P, Mallonee AB, Bergman NH, Koren S, Phillippy AM. 2016. Mash: fast genome and metagenome distance estimation using MinHash. Genome Biol 17:132.

59. Joensen KG, Tetzschner AM, Iguchi A, Aarestrup FM, Scheutz F. 2015. Rapid and easy in silico serotyping of Escherichia coli isolates by use of whole-genome sequencing data. Journal of clinical microbiology 53:2410–2426.

60. Larsen MV, Cosentino S, Rasmussen S, Friis C, Hasman H, Marvig RL, Jelsbak L, Sicheritz-Pontén T, Ussery DW, Aarestrup FM. 2012. Multilocus sequence typing of total-genome-sequenced bacteria. Journal of clinical microbiology 50:1355–1361.

61. Beghain J, Bridier-Nahmias A, Le Nagard H, Denamur E, Clermont O. 2018. ClermonTyping: an easy-to-use and accurate in silico method for Escherichia genus strain phylotyping. Microbial Genomics 4.

62. Siguier P, Perochon J, Lestrade L, Mahillon J, Chandler M. 2006. ISfinder: the reference centre for bacterial insertion sequences. Nucleic Acids Res 34:D32–6.

63. Clausen PTLC, Zankari E, Aarestrup FM, Lund O. 2016. Benchmarking of methods for identification of antimicrobial resistance genes in bacterial whole genome data. Journal of Antimicrobial Chemotherapy 71:2484–2488.

64. Clausen PTLC, Aarestrup FM, Lund O. 2018. Rapid and precise alignment of raw reads against redundant databases with KMA. BMC Bioinformatics 19.

65. Robertson J, Nash JHE. 2018. MOB-suite: software tools for clustering, reconstruction and typing of plasmids from draft assemblies. Microbial Genomics 4.

66. Jain C, Rodriguez-R LM, Phillippy AM, Konstantinidis KT, Aluru S. 2018. High throughput ANI analysis of 90K prokaryotic genomes reveals clear species boundaries. Nature Communications 9.

67. Gascuel O. 1997. BIONJ: an improved version of the NJ algorithm based on a simple model of sequence data. Molecular Biology and Evolution 14:685–695.

68. Paradis E, Schliep K. 2019. ape 5.0: an environment for modern phylogenetics and evolutionary analyses in R. Bioinformatics 35:526–528.

69. Guy L, Roat Kultima J, Andersson SGE. 2010. genoPlotR: comparative gene and genome visualization in R. Bioinformatics 26:2334–2335.

70. Hanson B, Johnson J, Leopold, Sr., Sodergren E, Weinstock G. 2019. SVants – A long-read based method for structural variation detection in bacterial genomes doi:10.1101/822312. Cold Spring Harbor Laboratory.

