## Supplemental Figures for "Systematic Analysis of Mobile Genetic Elements Mediating β-lactamase Gene Amplification in Non-Carbapenemase-Producing Carbapenem Resistant *Enterobacterales* Bloodstream Infections"

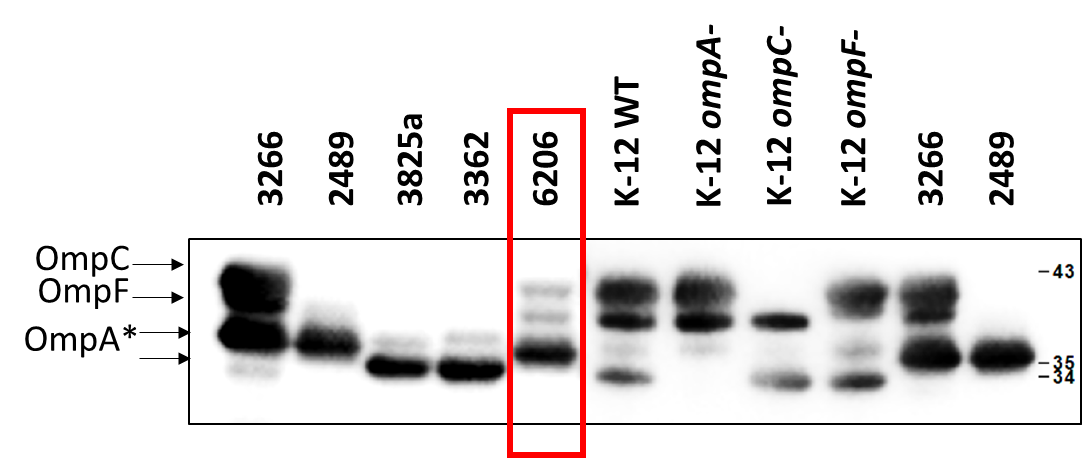

**Supplemental Figure 1. Immunoblot analysis** **of OmpC/OmpF protein levels.** Isolate MB6206 (red box) with IS*Ecp1*-*bla*_CTX-M-55_ insertion into the *envZ* gene displays reduced OmpC/OmpF levels as compared to clinical and lab positive control (MB3266 and K-12 WT, respectively). A series of clinical (MB2489,3825a,3362) and K-12 derived *omp* gene knockouts were used to aid with the band identification. *Note: MB3266, 2489, and 6206 contain a 4 amino acid insertion in OmpA protein as compared to K-12, resulting in a change of migration pattern. Immunoblot analysis was described before (1). Briefly, cell lysates were prepared from exponential phase cultures, grown in Lysogenic Broth. Samples were normalized by OD600. Proteins were separated on SDS polyacrylamide gels supplemented with 4 M urea. Immunoblots were developed with previously validated polyclonal rabbit antibodies raised against OmpA, OmpC, and OmpF (2, 3), and visualized using the ChemiDoc MP Imaging System (Bio-Rad).

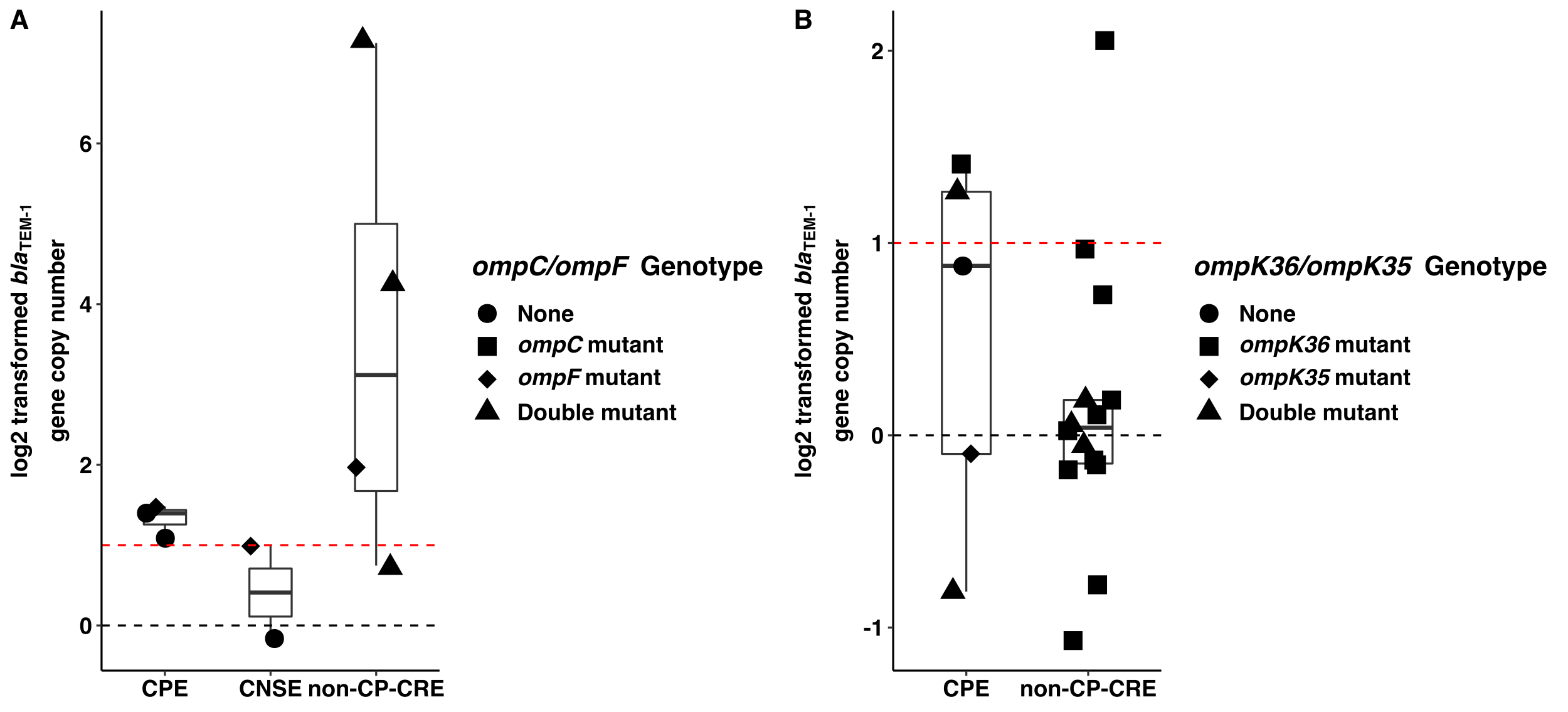

**Supplemental Figure 2. *bla*_TEM-1_ Log2 transformed copy number estimates for (A) *E. coli* and (B) *K. pneumoniae* stratified by CNSE group.** Black dotted horizontal line at y=0 is equivalent to 1X gene copy; Red dotted horizontal line at y=1 is equivalent to 2X gene copy.

**Supplemental Figure 3. Full Length Linear Comparisons of IncF type plasmids.** Transposases/integrases (dark grey), IS*26* transposase (white), IS*26*-v1 (off-white), IS*Ecp1* transposase (purple), carbapenemases (orange), other antimicrobial genes (red), *rep* genes (yellow), *bla*_OXA-1_ (blue arrow), *bla*_CTX-M-15_ (green arrow), virulence factors (pink), and other genes (light grey) are labelled accordingly. Striped IS*Ecp1* transposase ORFs indicate a disruption due to IS*26* or IS*26*-v1). Linear comparisons between sequences indicate homology shared (min length 1000 bp and 90% identity) in direct (red) and reverse (blue) orientation.

**REFERENCES**

1.	Tata M, Konovalova A. 2019. Improper Coordination of BamA and BamD Results in Bam Complex Jamming by a Lipoprotein Substrate. mBio 10.

2.	Misra R, Reeves P. 1987. Role of micF in the tolC-mediated regulation of OmpF, a major outer membrane protein of Escherichia coli K-12. Journal of bacteriology 169:4722-4730.

3.	Zimmermann R, Wickner W. 1983. Energetics and intermediates of the assembly of Protein OmpA into the outer membrane of Escherichia coli. Journal of Biological Chemistry 258:3920-3925.
